# Converting cropping systems into seasonal habitat functionality reveals the hump-shaped responses of carabid beetles to agricultural management

**DOI:** 10.1101/2022.02.26.480658

**Authors:** Lucile Muneret, Benoit Ricci, Aude Vialatte, Stéphanie Aviron, Chantal Ducourtieux, Luc Biju-Duval, Sandrine Petit

## Abstract

1. Understanding effects on the huge diversity of cropping systems on local biodiversity is challenging but necessary to implement agroecological systems. Through a functional approach, the translation of cropping systems into resource and disturbance gradients is promising to decipher the relationship between cropping systems and biodiversity but has never been implemented for arthropods.
2. To investigate contributions of resource and disturbance gradients arising from cropping systems *vs* environmental context (regional effect, meteorological conditions and landscape characteristics) on beneficial arthropod communities, we used a dataset collected in 60 crop fields from three French areas over a five-years period. It includes all farmers interventions, crop sequences, meteorological data, landscape composition and carabid samplings.
3. We found that the environmental context contributed to about 75% of explained carabid variations on average, while resource and disturbance gradients contributed to about 25% of explained carabid variations. The resource and disturbance gradients were particularly important in winter and spring preceding the spring-summer period to determine carabid variations.
4. Moreover, we identified thresholds above which resource and disturbance gradients start being beneficial or detrimental for carabids. For example, a Treatment Frequency Index above 2.07 in spring decreased the total activity density of carabids during the spring-summer period.
5. Synthesis and application. While implementing for the first time a functional approach to understand the effects of different facets of cropping systems on arthropods, our study also allows us to identify periods and thresholds above which specific practices affect carabids. The identification of such thresholds can guide the provision of recommendations for policy, stakeholders and farmers about how to reduce cropping systems’ impact on arthropods.

## 1 INTRODUCTION

Mainstream agriculture is often associated with a decline of in-field biodiversity (Uhler et al., 2021). The expansion of alternative systems, such as organic farming or conservation agriculture may alleviate this trend (Petit et al., 2020). However, our understanding of how specific cropping systems, i.e. the crop sequence and the logical suite of farming operations implemented over successive years, affect different biodiversity components is still limited (Shennan et al., 2017). One aspect complex to tackle is that, even within broad types of farming systems, there is a huge diversity of practices and this so-called ‘hidden heterogeneity’ is a strong driver of biodiversity responses (Vasseur et al., 2013). Improving how we account for cropping systems to describe habitat functionality and how it varies over time thus remains challenging (Schellhorn et al., 2015), but has the potential to increase our capacity to predict biodiversity trends in agroecosystems.

Some conceptual frameworks have emerged postulating that crop sequences and associated farming operations alter the amount and timing of resources available to or disturbances faced by species (Damour et al., 2017). They thus propose to translate cropping systems into ecological gradients of resource availability and disturbance intensity (Gaba et al., 2014). This framework was primarily developed for plants and successfully implemented to describe crop sequences along gradients of disturbance and resource variability for weeds (Mahaut et al., 2019). It has never been applied to arthropods in agroecosystems, whereas these organisms are likely to respond to both gradients. This represents a major gap in agroecological studies as arthropod species provide key ecosystem services like biological control (e.g. Gagic et al., 2017).

Among agrobiont arthropods, carabid beetles are the most abundant group in temperate arable agriculture. Carabids consume many prey types and are involved in pest and weed controls (Frei et al., 2019). Effects of agricultural management on carabids has been widely documented, either broadly, through comparisons of pairs of fields grown with the same crop but under contrasting farming systems, e.g. organic vs. conventional (e.g. Diekötter et al., 2010) or, very specifically, through experimentation, mostly to quantify the impact of a single farming practice e.g. tillage intensity (Shearin et al., 2007), fertilisation type (Aguilera et al. 2021) or pesticide use intensity (Geiger et al., 2010). Some studies described local habitat quality for carabids without using information on farming practices, for example with estimates of in-field available trophic resources (Carbonne et al., 2022) or proxies for habitat productivity and disturbance (Eyre et al., 2013). This body of literature yielded mixed results; probably because 1) they only tested simple linear responses of carabids to variations of habitat conditions while their responses could have quadratic shapes and 2) they did not account for the sheer diversity of cropping systems found in arable landscapes. Investigating such knowledge gaps could help to understand carabid responses to management intensity.

Another aspect requiring attention is the timing of farming operations, and how it may interfere with different carabid life stages. Carabid responses to environmental factors are generally examined via snapshots of both the community and environmental conditions (Schellhorn et al., 2015). However, environmental conditions, as well as carabid species requirements vary over a cropping season. Accounting for the timing of farming operations is thus key. Fields are expected to supply to carabids with different life-cycles, trophic resources during the growing or active period, partners during the breeding period and overwintering sites during winter. Some practices implemented at different periods can have different impacts on species or community composition (Timberlake et al., 2019; Marrec et al., 2021). This calls for a seasonal account of the habitat functionality created by cropping systems.

In addition to cropping systems, carabids also respond to the environmental context, notably to landscape and climatic variations. The landscape context of arable fields is known to affect carabids, with often a higher species richness in complex landscapes (Purtauf et al, 2005) but less consistent results for carabid abundance (Winquist et al., 2011; Djoudi et al., 2019). Furthermore, temperature modulates the presence or activity of individuals, with an effect on the number of catches in pitfall traps (Honek, 1997; Eyre et al., 2005). However, the effect of meteorological conditions on carabid distributions has extremely rarely been considered while such variables could be very important to increase our understanding.

Here, we characterised the ecological conditions created by cropping systems and tested their relevance to explain carabid distribution during the spring-summer period, when carabids contribute to pest control. We developed an approach in which technical routes are translated into two ecological gradients, i.e. resource availability and disturbance intensity. Data came from 60 arable fields located in three French regions and monitored from 2014 to 2018, covering 31 crop species. Our first objective was to quantify the influence of resource and disturbance gradients created by cropping systems on carabid distribution and compare this effect to that of “environmental context”, meaning here variations of region, meteorological conditions and landscape context in space and time. We expected that fields providing a high amount of resources and limited disturbance would be highly suitable for carabids. Moreover, we hypothesized that the relationship may not be linear but rather hump-backed, with some optimal values of resources and disturbance for carabids. Our second objective was to identify periods over the year during which these ecological gradients affect carabid distribution in the spring-summer period. We expected seasonal factors to be better predictors than annual factors. Specifically, we expected a maximal positive effect of resource availability in spring when carabids emerge. In contrast, we expected a maximal negative effect of the physical disturbance intensity during autumn and winter when carabids overwinter and the maximal negative effects of chemical disturbances in spring and autumn when carabids circulate.

## 2 METHOD

### 2.1 Sites and carabid community description

Sixty arable fields were selected according to two crossed gradients, the proportion of cultivated habitat in the landscape (1km^2^) and local agricultural management intensity (French Landscape Monitoring Network SEBIOPAG; Petit et al., 2021). We annually monitored 20 fields in three French regions (near Dijon 47.32N, 5.04E, Rennes 48.12N, -1.68E and Toulouse 43.60, 1.44; Figure 1) between 2014 and 2018. Crop rotations over the 5-years period included 31 crop species (Table S1). Managed by 52 farmers, 11 fields were in conservation agriculture, 13 in organic farming, and 36 under conventional agriculture with contrasted strategies.

**Figure 1.**
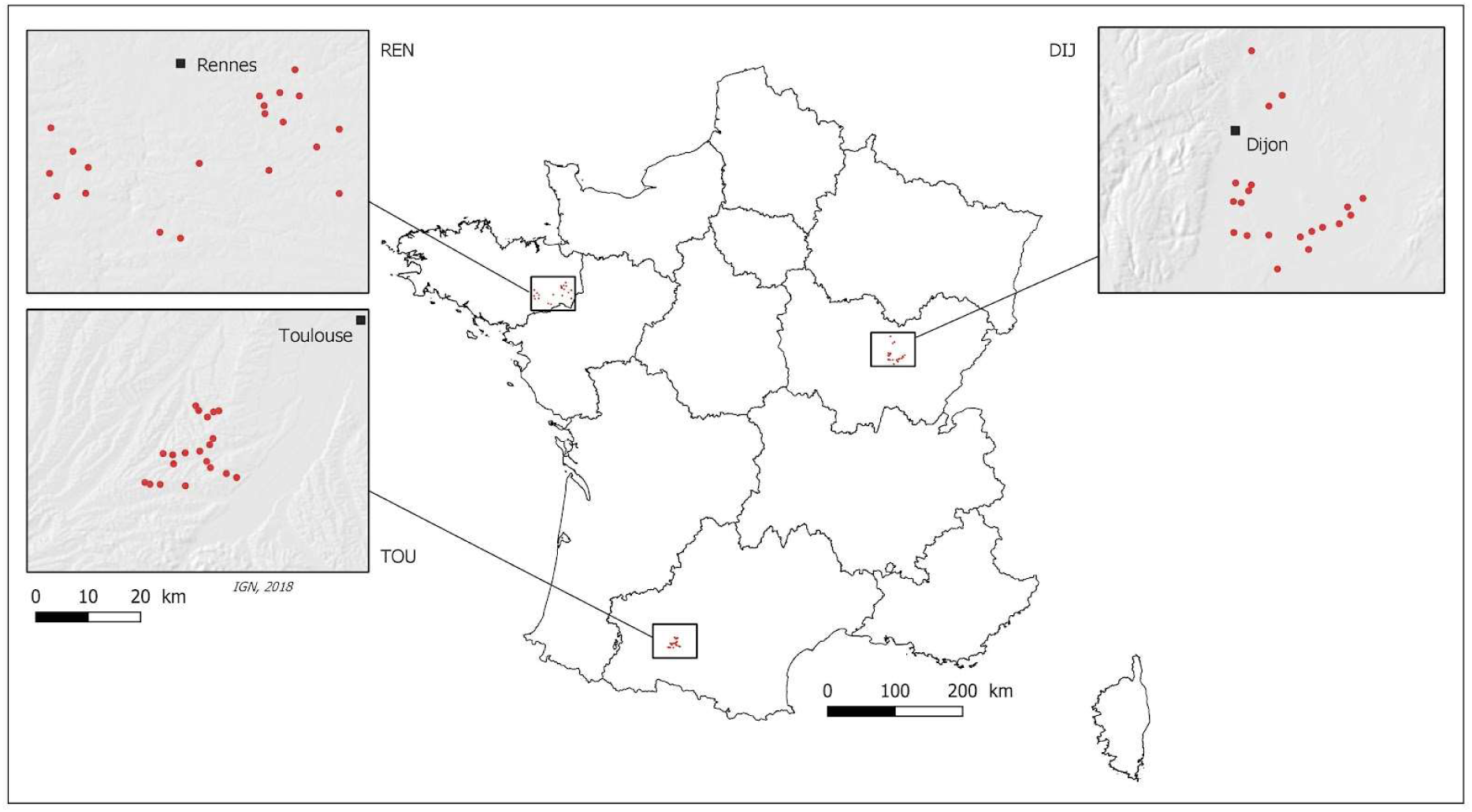
Map of the study sites.

Carabids were sampled within fields by opening four pitfall traps (10 cm long * 8 cm diameter) on the ground over four days twice a year, from 2014 to 2018. Two pitfall traps were placed at 10 m from the edge and each other. The two other traps were at 50 m from the edge. To make carabid community samplings comparable over latitudes, sampling dates were determined according to degree days. They were conducted at 1100 and 1500 degree-days for winter crops. For summer crops, the former sampling took place at 1500 degree-day and the latter occurred after about the same interval of time that there was between the two dates of winter crop samplings. Specimens were identified at the species level except some rare specimens (genus level).

The community of each field and sampling date was described with two metrics, namely species richness and activity-density. We also tested our conceptual framework at a species level (Jowett et al., 2019), on the occurrence of the six most abundant species collected in the three regions (Figure S1). *Poecilus cupreus* and *Brachinus crepitans* (generalist species), *Anchomenus dorsalis* and *Trechus gr. quadristriatus* (hereafter ‘*T. quadristriatus*’) are carnivorous species, *Pseudoophonus rufipes* and *Amara similata* are phytophagous species. We used species occurrence as a response variable to account for behavioral differences between species (agrarian or not).

### 2.3. Translating cropping systems into ecological gradients

Through interviews, farmers provided all the interventions from their technical routes with dates, agricultural machineries, products and applied doses for every cash and cover crops. The first day of this 5-year sequence was 12 August 2013 and the last day was the last harvest in 2018. We converted all the interventions into two ecological gradients: resource availability and disturbance intensity (Figure 2; Figure S2)

**Figure 2.**
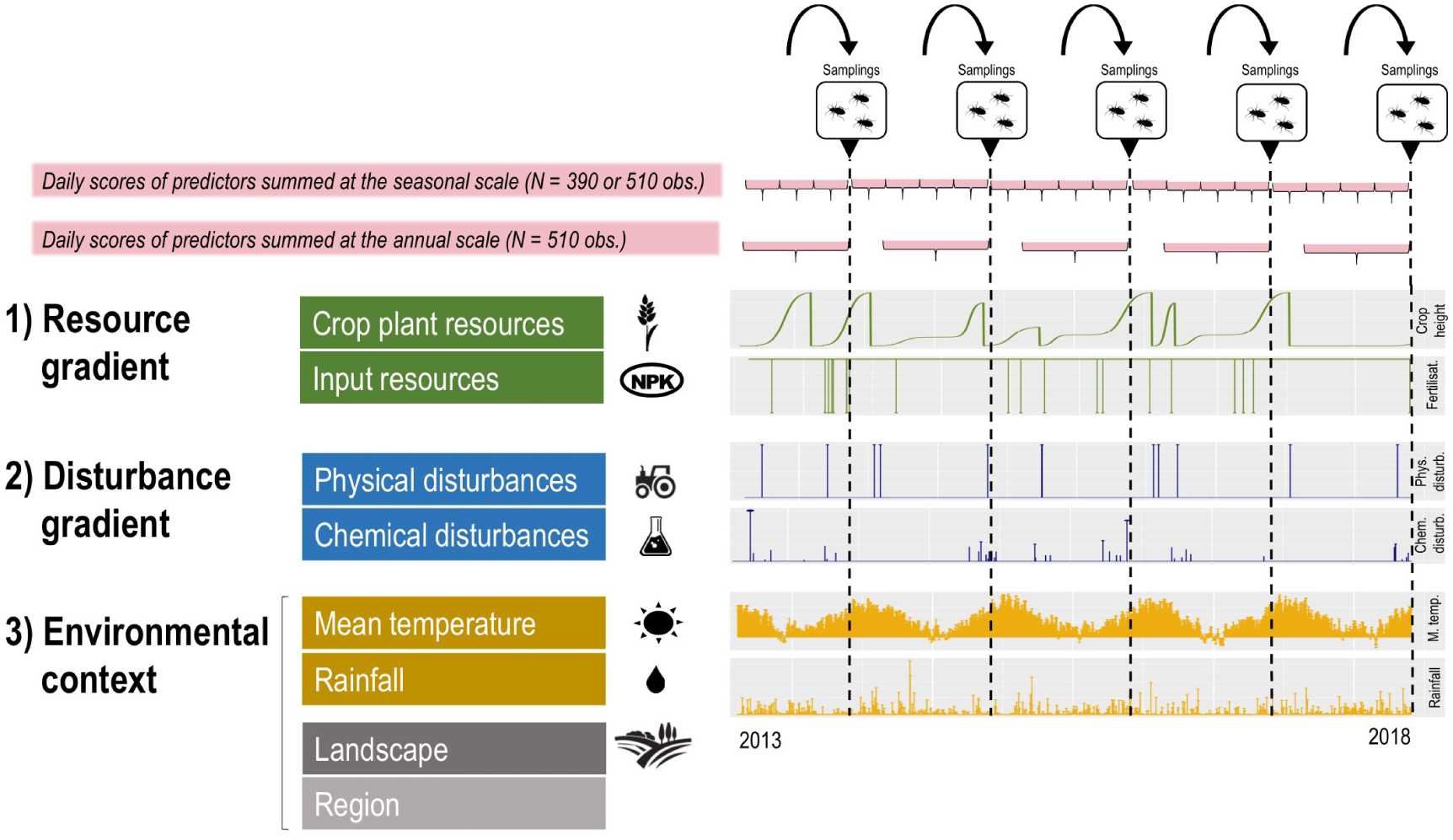
Calculation of environmental variables according to the conceptual framework.

#### 2.3.1 Crop plant resource

We assumed that crop biomass was a driver of carabid communities as it was shown to increase arthropod biomass at multiple trophic levels (Fernández-Tizón et al., 2020). We selected crop height as a proxy of resource availability because it is positively correlated with crop biomass (e.g. Yin et al., 2012; Bendig et al., 2015). Based on the maximal crop height extracted from the database LEDA (Kleyer et al., 2008) or agronomist appraisals, sowing and harvest dates, we calculated an estimated crop height each day over the 5-years period in each field. A few missing dates on sowing or harvest were estimated with sowing and harvest dates from other fields in the same area. Three typologies of crop growth were calculated: logistic growth, logistic crop growth with a break in winter and grazing (Annexe 1).

#### 2.3.2 Farming inputs resource

We assumed that organic matter inputs provide prey for carabids while mineral application can reduce their availability (Riggi et Bommarco, 2019). Because of the lack of consensus on the effect of subsidy inputs on biodiversity, i.e. including potential resources for carabid communities (Tamburini et al., 2020), we applied a qualitative approach. One organic input has a positive effect on carabid resources and a mineral application has a negative effect on carabid resources (score of organic fertilisation set to one, score of mineral fertilisation set to minus one).

#### 2.3.3 Chemical disturbance

The highest scores of chemical disturbances mean higher levels of pesticide application. We expected that low levels of pesticide applications have weak effects on carabids while higher levels should be very detrimental to carabids. We estimated the level of chemical disturbance for each pesticide application using the Treatment Frequency Index “TFI”

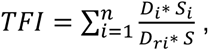

where *D_i_* is the applied dose, *S_i_* the treated surface area, *D_ri_* the reference dose obtained from the French Ministry of Agriculture database (Ephy website) and *S* the total area of the field for each spraying operation *i*. All the TFI were added to have a daily score of chemical disturbance intensity.

#### 2.3.4 Physical disturbance

The highest scores of physical disturbances were expected to be much more detrimental to carabids than the lowest levels of physical disturbances. Based on agronomist appraisals, we ranked every farmer’s operations on a scale relative to the magnitude of its negative effect on physical habitat stability. The more in-depth and animated, the more disturbing (Table S2). When two different agricultural machineries were employed the same day, their scores were added to provide a daily score.

### 2.4 Environmental context

Daily meteorological data came from three meteorological stations close to sampling plots in each region (CLIMATIK database). Variables were daily mean temperature (degree celsius) and rainfall (mm). Over the 5-years period, 51 days of rainfall and seven days of mean temperature were missing. We estimated the values on those days with the mean of the given variable calculated on data from the four other years for the given region. We obtained one measure a day for both meteorological variables in each field.

To describe landscape contexts, land covers were characterised in a 1 km^2^ buffer circle around focal fields using the ALM procedure each year (Allart et al., 2021). The proportion of cultivated area was calculated as an indicator of land use intensification. Arable and perennial crops were included in the cultivated area except permanent meadows, classified as semi-natural habitats.

### 2.5 Calculation of annual and seasonal explanatory variables

Explanatory variables estimated each day over the 5-years period were aggregated at two levels of resolution, annual and seasonal levels. We considered four seasons: summer (June - August), autumn (September - November), winter (December - February) and spring (March - May). Annual periods were considered between 1 September and 31 May because data missed for the summer period in 2013. For each environmental variable, we summed daily scores for each period.

Explanatory variables describing the ecological gradients created by the cropping systems were standardised at the scale of the whole dataset, like the landscape metric. This allowed us to compare parameter estimates with each other. We assumed that practice’ choice varied according to farmers’ strategies, independently of regions. Variations of resource and disturbance gradients relative to seasons were evaluated using gaussian linear models and Tukey tests. Conversely, meteorological variables were standardised at the regional scale. With this, we evaluated the effect of annual variation of such variables on the period at the regional scale. We hypothesised that carabids were in the range of distribution and well adapted to each region and that annual meteorological variations influence carabid activity density.

### 2.6 Modelling

To evaluate the effects of seasonal and annual variations of environmental variables on carabid distribution, we developed eight sets of mixed effects models, i.e. one for each response variable. Sets were composed of five models, one for each period (seasons and year). For species richness and activity density responses, we developed negative binomial mixed effects models and binomial mixed effect models for the occurrence of species. Explanatory variables were “region”, “proportion of cultivated area”, “mean temperature”, “rainfall”, “crop height” and “fertilisation”, “physical disturbance” and “chemical disturbance”. Quadratic effects of all the quantitative explanatory variables were also included. One random term “field” and an offset describing the number of used pitfall traps per field were also added in the models. Collinearity between explanatory variables has been previously checked (Figure S3). Annual, autumn, winter and spring models were fitted with 510 observations and summer models with 390 observations.

We applied a multimodel inference approach to evaluate the effect of explanatory variables on response variables (Burnham et al., 2011). All the possible models were ranked according to their AIC and models with a ΔAIC < 2 were retained in the set of top models. Using model averaging, top models were then used to calculate parameter estimates and their confidence interval. We calculated marginal and conditional R^2^ based on the model with the lowest AIC (function “r.squaredGLMM”; package “MuMIn”; Bartoń, 2020). As we developed multiple models, we adjusted p-values with the “false discovery rate” method (Benjamini et al., 2001). Finally, to calculate the contribution of the explanatory variables to response variations, we summed the absolute values of parameter estimates from the “average model” and divided the sum of absolute parameter estimates of the compartment by the total sum of the parameter estimates for this model. It corresponds to variance partitioning because we previously scaled all the explanatory variables. All the models were performed with R (R Core Team, 2021), “lme4” (Bates et al., 2015) and residuals were checked with “DHARMa” (Hartig, 2021).

For each predictor retained by the model averaging procedure, we calculated the turning point (explicitly given by the formula *-b/2c*) associated with a quadratic response (*y=a+bx+cx^2^*) by finding the root of the derivative of the quadratic function. Here, *y* is the response variable with respect to the predictor *x*, *a* the intercept, *b* and *c* are the first and second-order of the predictor (Annexe 2). Negative value of *c* means that the predictor has a hump-backed effect on the response while a positive value is associated with an inverse bell-shaped effect. To observe such effect, the turning point has to be included in the range of values of the predictor. Otherwise, the effect only has a positive or negative effect in the range of values of the predictor (Annexe 2). The magnitude of *c* reveals the magnitude of the quadratic effect.

## 3 RESULTS

Our analyses were based on a collection of 19839 carabids from 116 taxa. The three regions had different temporal patterns of activity density (Figure S4). Resource and disturbance gradients, substantially varied according to seasons (Annexe 3). Furthermore, we obtained a value of agricultural intensification per field each year ranging from 8% to 99% of cultivated area in the 1km^2^ buffer (Figure S5) and meteorological conditions varied according to regions and periods. The results and discussion of specific effects of the environmental context on carabid distributions are presented in Annexe 4.

### 3.1 Effect of resource gradients on carabid distribution

On average, resource availability accounted for about 14% of explained variance of carabid distributions in the set of best models retained (Figure 3). The contribution of resource availability to explained variance ranged from about 6% for *H. rufipes* to about 40% for *A. similata* (Table S3). Resource availability in spring and winter was more important to explain carabid distribution (25 and 15%) than over other periods (from 6% to 12%). Crop height accounted for 11% of the explained variance while fertilisation accounted for 3% on average (Figure 3).

**Figure 3.**
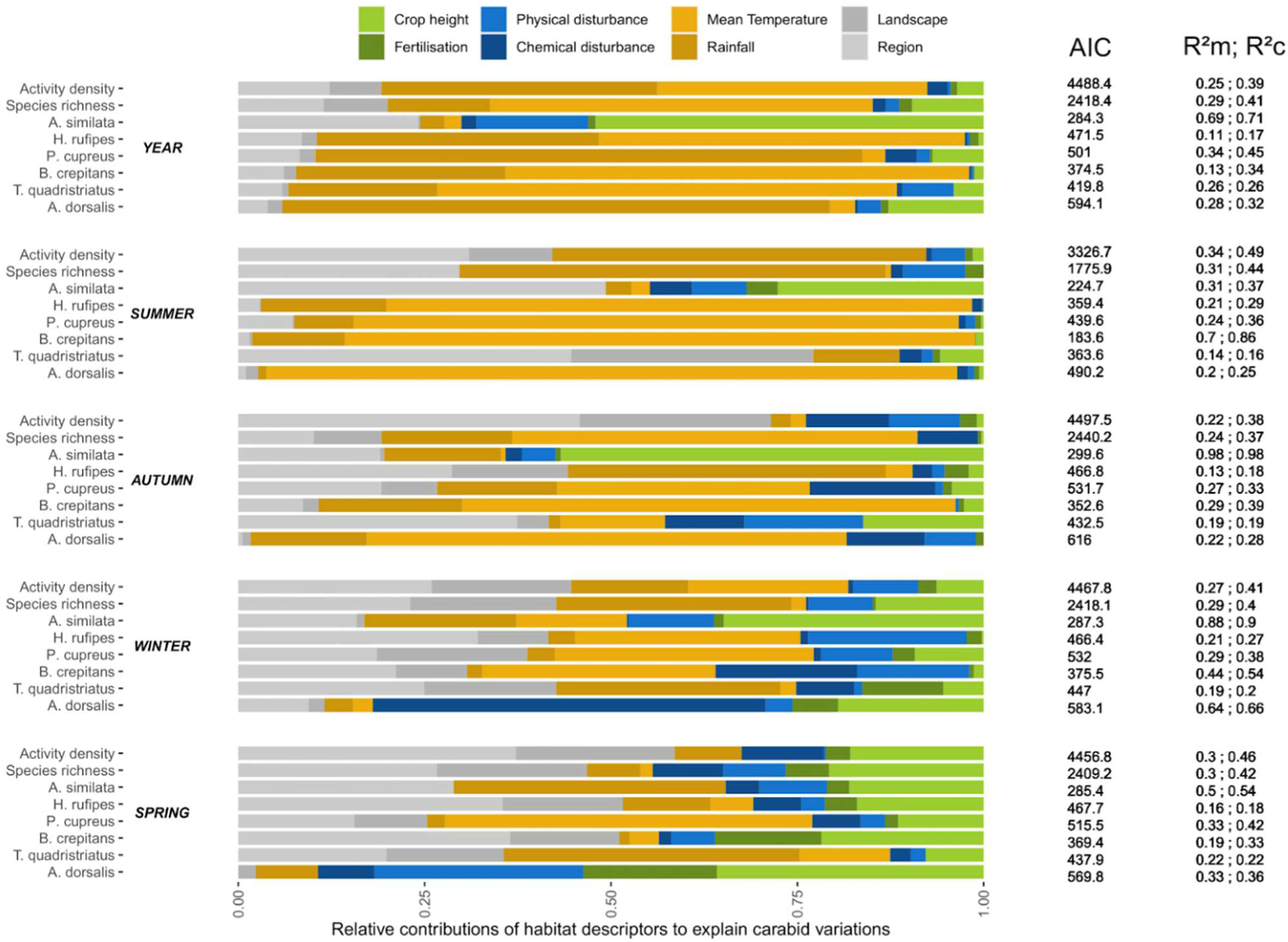
Relative contribution of explanatory variables to explain carabid distributions. Crop height and fertilisation are predictors describing the resource gradient; chemical and physical disturbances describe the disturbance gradient; mean temperature, rainfall, landscape and region describe the environmental context

Crop height had a significant effect on carabid distribution in 21 out of 40 models tested and this effect was consistently hump-shaped (Figure 4) except in five cases. *A. similata* had a positive response to crop height in annual, autumn and winter models, as *H. rufipes* in summer models while *A. dorsalis* had an inverse bell-shaped response in summer models. Crop height had their main effect on carabids in spring models (19% of explained variance on average).

**Figure 4.**
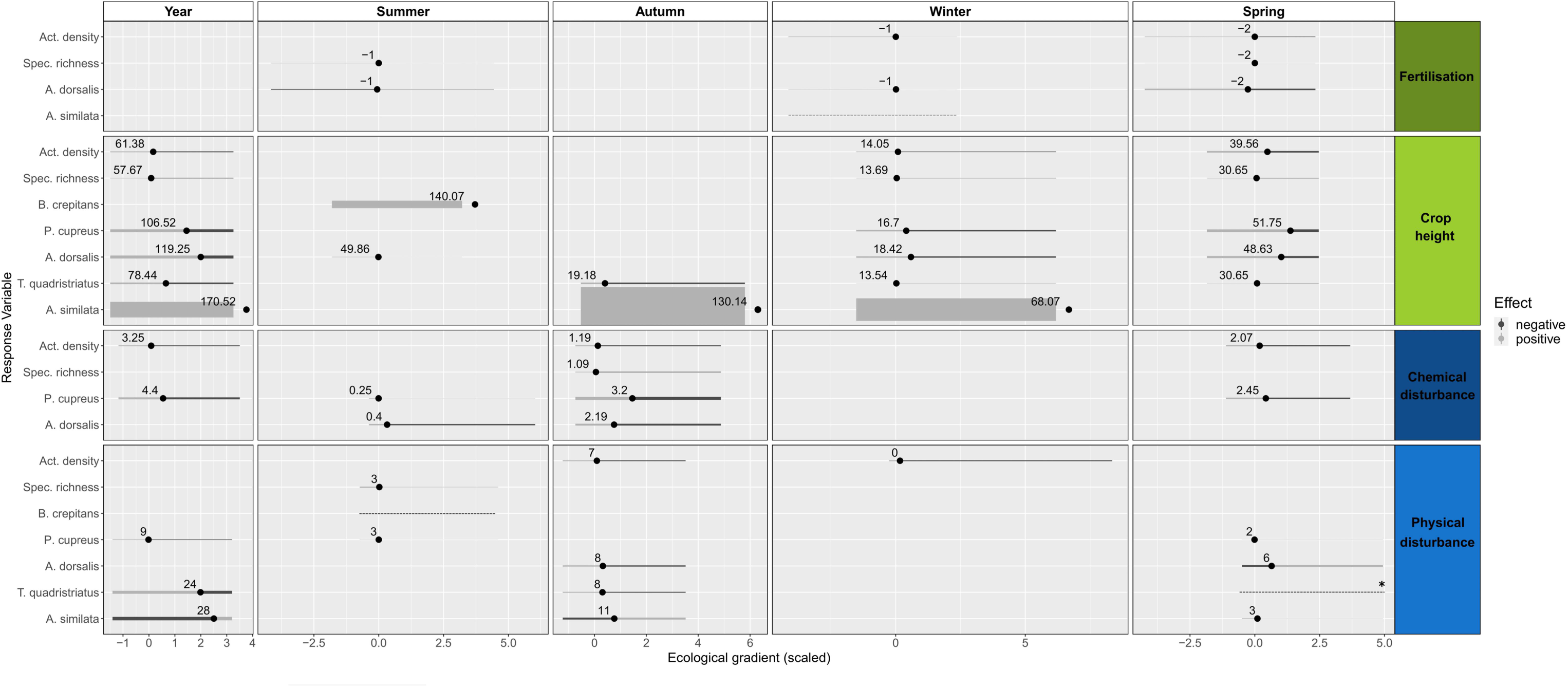
Carabid responses to resource and disturbance gradients. Two-tone rights represent quadratic effects with a negative slope for the blue portion and a positive slope for the red one. One-tone right displays simple linear effects. Right thicknesses represent the intensity of the effect and are proportional to the magnitude the parameter *c* of each predictor. Only significant effects are represented (Table S3). Rights without dot have only simple linear effect. For *T. quadristriatus* (spring, physical disturbance), only quadratic effects have been retained in the set of best models, this effect is therefore complicated to interpret.

Fertilisation had a significant effect on carabid distribution in eight out of 40 models (6% of explained variance on average; Figure 3). Its effect greatly varied according to periods. In summer models, species richness and occurrence of *A. dorsalis* responded positively to high levels of both mineral and organic fertilisations. The same pattern was detected for activity density and the occurrence of *A. dorsalis* in winter models. Conversely, in spring, activity density, richness and occurrence of *A. dorsalis* were negatively affected by high levels of fertilisation, here above two mineral applications.

### 3.3 Effects of disturbance gradients on carabid distribution

On average, disturbance gradients accounted for about 11% of the explained variance of carabid distributions and were the most influencing in winter and spring models (20% and 14%). Both disturbance gradients, i.e. chemical and physical, accounted each for around 6% of explained variance on average (Figure 3).

The effect of chemical disturbance was significant in 10 out of the 40 models, with very consistent bell-shaped curves in 9 out of 10 models (Figure 4; Table S3). The threshold above which the chemical disturbance gradient had a negative effect on carabid varied from 0.4 to 3.2 of TFI points for seasonal models.

The physical disturbance gradient had a significant effect on carabids in 15 of the 40 models (Table S3). In half of the cases, we detected a bell-shaped response with low levels of physical disturbance benefitting carabids while the highest levels of physical disturbance having a negative effect on carabids. For such models, the range of thresholds until which the level of physical disturbance benefitted carabids varied from 0 to 8 points for seasonal models and from 9 to 24 points of or annual models. In six models, carabid responded to the physical disturbance gradient with an inverse bell shape, suggesting that very low and very high levels of physical disturbance intensities benefitted carabids.

### 3.4 Seasonal *vs* annual variations of environmental variables

Based on comparisons of annual, autumn, winter and spring models AIC for every response variable, models developed using seasonal predictors for carabid activity density, species richness, *A. dorsalis*, *B. crepitans, H. rufipes* performed better than those based on annual predictors (Figure 3; Table S3). For activity density, species richness, *A. dorsalis and B. crepitans* occurrences, models fitted with spring variables had the lower AIC while winter models had the lowest AIC for *H. rufipes*. Conversely, annual models had the lowest AIC for models based on *A. similata*, *P. cupreus* and *T. quadristriatus*.

## 4 Discussion

We implemented a functional and generic approach to explain arthropod distributions within agroecosystems. With this conceptual framework, we evaluated effects of a wide array of cropping systems including 31 crop species, on carabid distributions through their conversion into resource and disturbance gradients. Although we found that meteorological conditions were a surprisingly important driver of carabid distribution (46% of explained variation), we demonstrated that the ecological gradients created by cropping systems also shaped carabid distribution (25% of explained variation). We showed that the majority of community and species responses to resource and disturbance gradients were bell-shaped, highlighting optimums of resource availability and disturbance levels for carabids. We also identified which periods of the year carabids are the most sensitive to resource and disturbance gradients meaning winter and spring but that effects can greatly vary according to response variables.

### 4.1 Relative effects of environmental context *vs* resource and disturbance gradients

Variables describing the environmental context are the most important drivers of carabid distributions as they contributed to about 75% of explained carabid variations on average (respectively 46%, 8% and 19% of contributions for meteorological, landscape and regional variables). Relatively, resource and disturbance gradients contributed to about 25% of explained carabid variations. To our knowledge, previous studies did not include meteorological conditions as explanatory variables in their models, we therefore did not find comparative studies. In contrast, the effect of landscape composition on arthropods has been massively investigated but is still ambiguous. It is highly context-dependent (Karp et al., 2018) because most of studies implicitly include meteorological conditions as well as cropping systems within the landscape variation while we unravelled all these effects here.

In comparison to the environmental context, resource and disturbance gradients, representative of the cropping systems, were rarely integrated together within unique studies. Generally, they are included as farming systems, proxy of vegetation cover or even effects of specific practices are evaluated. Organic farming or conservation agriculture are considered as positive for carabids (e.g. Henneron et al., 2015) and the effect of single practices depends on studies (Jowett et al., 2021; Shearin et al., 2007). The advantage of our approach is that we unraveled the relative contribution of the different practices being part of coherent cropping systems (Damour et al., 2017). Our results suggest that farmers’ capacity to increase in-field carabid activity through change of the technical routes is possible but have not massive consequences. This suggests that such changes must be accompanied by changes at wider scales, notably at the landscape scale or even in a way that contributes to mitigate climate change. These results coincide with studies showing that landscape-level crop diversity enhance pest control (Redlich et al., 2018) and that some specific crop rotations considered at the landscape scale greatly influence carabid distributions within agricultural landscapes (Marrec et al., 2017).

### 4.2 Bell-shaped effects of resource gradients on carabids

Crop height and fertiliser applications had bell-shaped effects on carabid variations in most cases. This shows that a minimum of resources is necessary for carabid activity but that too high levels of resources reduce their activity.

Beyond the well-known effect of crop identity on carabids (Jowett et al., 2021), we showed that crop height, especially in winter and spring, drives carabid distributions independently of its identity. The effect may arise because crop height is correlated with crop biomass (e.g. Yin et al., 2012) that can have a positive effect on arthropod biomass consumed by carabids (Fernandez-Tizon et al., 2020). The general bell-shaped effect of crops in annual, winter and spring models can result from several mechanisms. One the one hand, maintaining cover crops during winter could increase the capacity of fields to shelter overwintering carabids (Dennis et al., 1994). Moreover, intermediate crop height in spring may mean that carabids find prey when they emerge, which favours their survival. On the other hand, it is possible that the highest crop height limits carabids because the recruitment of other species like spiders could limit them via intra-guild predation (e.g. Roubinet et al., 2018). Another explanation is that carabid movements could be limited by the development of plants and that carabids could be more competitive in simpler habitats. In comparison, fertiliser applications often had a weak effect on carabids, either negative or positive effects depending on their type (synthetic or organic) and on periods. This result confirms the lack of consensus on the effect of fertilisers on biodiversity (Tamburini et al., 2020).

### 4.3 Contrasting responses of carabids to physical and chemical disturbance gradients

Contrary to our hypothesis, carabid response to increased levels of disturbance was not homogeneous. Effects depended on the type of disturbance and, for physical disturbances, on species and periods.

We consistently detected a bell-shaped response to chemical disturbances (except once) and identified thresholds for each period above which pesticide applications start being detrimental to carabids, here 1.09 to 3.2 in autumn and 2.07 to 2.45 in spring, values depending on the carabid metrics. Although pesticide applications are detrimental to carabids and biodiversity in general (Geiger et al., 2010), our results identified for the first-time harmful thresholds for several carabid variable responses. Furthermore, several species did not respond to pesticide use intensity, suggesting that they might have developed resistance mechanisms to pesticides or they only experienced sublethal effects, with no effect on their abundance but an impact on their nutritional state (Labruyere et al., 2016).

Unlike the homogeneous response to the chemical disturbance gradient, the effect of the physical disturbance gradients on carabids depended on response variables and periods. This probably reflects the diversity of carabid responses to soil cultivation (Shearin et al., 2007; Ward et al., 2011) and likely the huge diversity of strategies encapsulated within the carabid community. It is possible that to enhance carabids through tillage strategies, reasoning at both the landscape or crop rotation scales could be relevant (Petit et al., 2020). A high diversity of tillage strategies at a landscape scale could thus enhance the occurrence of multiple abundant species.

### 4.4 Effect of the environmental context on carabids

The effect of environmental context contributed to about 75% of the explained of carabid variations. Meteorological conditions were the most influencing variable. On average, it explained about 46% of variations in carabid distributions while the proportion of cultivated area in the landscape accounted for 8% and region 20%. Extended discussion is presented in Annexe 4.

### 4.5 Are seasonal habitat patterns useful?

Decomposing the year by seasons was an asset because all the seasons did not have the same weight on the spring-summer carabid distributions. We showed that practices implemented during winter and spring were the most influential on carabids whereas meteorological conditions in summer had the highest influence. Another finding is that a specific farming operation does not have the same weight on carabids depending on its timing in the year. This result calls for further studies conducted at the intra-annual scale (Schellhorn et al., 2015), with potential to uncover mechanisms of community assembly (Marrec et al., 2021).

## 5 Conclusions

Implementing a functional approach to translate cropping systems into resource and disturbance gradients is very useful to understand which facets of such systems affects the in-field arthropod community. We showed that the environmental context drives the majority of the explained carabid variations within agroecosystems (about 75%). This supposes to include meteorological conditions in a more systematic way when analyses are conducted at wide temporal or spatial scales. Moreover, the conversion of cropping systems into resource and disturbance gradients contributed at an extent of about 25% of explained carabid variations. Specifically, these gradients were substantially influential in the winter and spring seasons preceding the spring-summer period. In addition, carabid responses were in most cases hump-shaped highlighting optimal conditions of resources and disturbances for the community and dominant species. The identification of such optimums, as thresholds, can guide the provision of recommendations for policy, stakeholders and farmers about how to reduce cropping systems’ impact on carabids. This approach needs to be extended to other functional groups to identify trade-offs between them and with ecosystem services. This could allow us to understand why some species are winners or losers in contrasting farming systems.

## Authors’ contributions

L.M., B.R., A.V., S.A., S.P. designed the study. C.D. and L.B.D. managed the data. L.M. and B.R analysed the data. L.M. wrote the first draft. S.P. acquired and managed funding. All authors interpreted the results, contributed to the drafts and gave final approval.

## Acknowledgements

We are grateful to Sylvie Ladet, Diane Esquerré, Jérôme Willm, Jérôme Molina, Laurent Raison et Bruno Dumora, Audrey Alignier, Alexandre Joannon, Jean-Luc Roger, Gérard Savary, Kristell Jegou, Ninon Frémaux, Arnaud Maillard; Manuel Plantegenest, Yann Laurent, for their technical help; Rolland Allart for calculating landscape metrics and the partner farmers. Funding came from the Ecophyto Plan of the French Ministry of Agriculture and Food and the French Foundation for Research on Biodiversity, projects ‘SEBIOPAG-PHYTO’ and ‘SEBIOPAG-STAR’.

## Data availability

https://doi.org/10.15454/BMLIQI

## Supplementary materiel

**Table S1.**
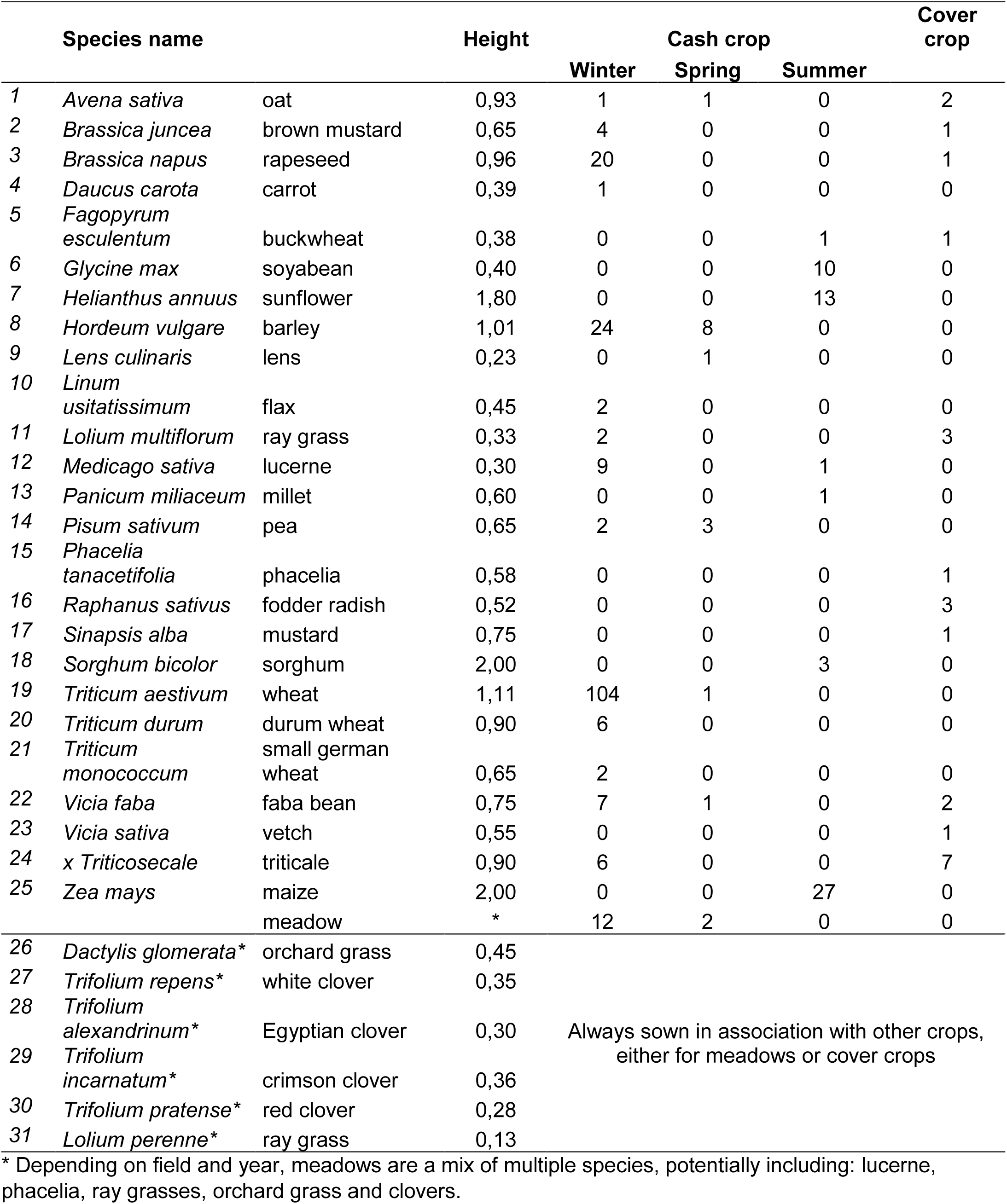
The 31 crop species grown in the surveyed fields between 2014 and 2018. Crop species were used in cash crops, and sometimes in cover crops. Some crop species could be used in crop mixtures. * indicates that the crop species was always sown in association with other crop species, either in meadows, cash or cover crops. Height is the maximal height of the highest leaf of the canopy at the ripening stage; values were extracted from the LEDA database or were expert-based.

**Table S2.**
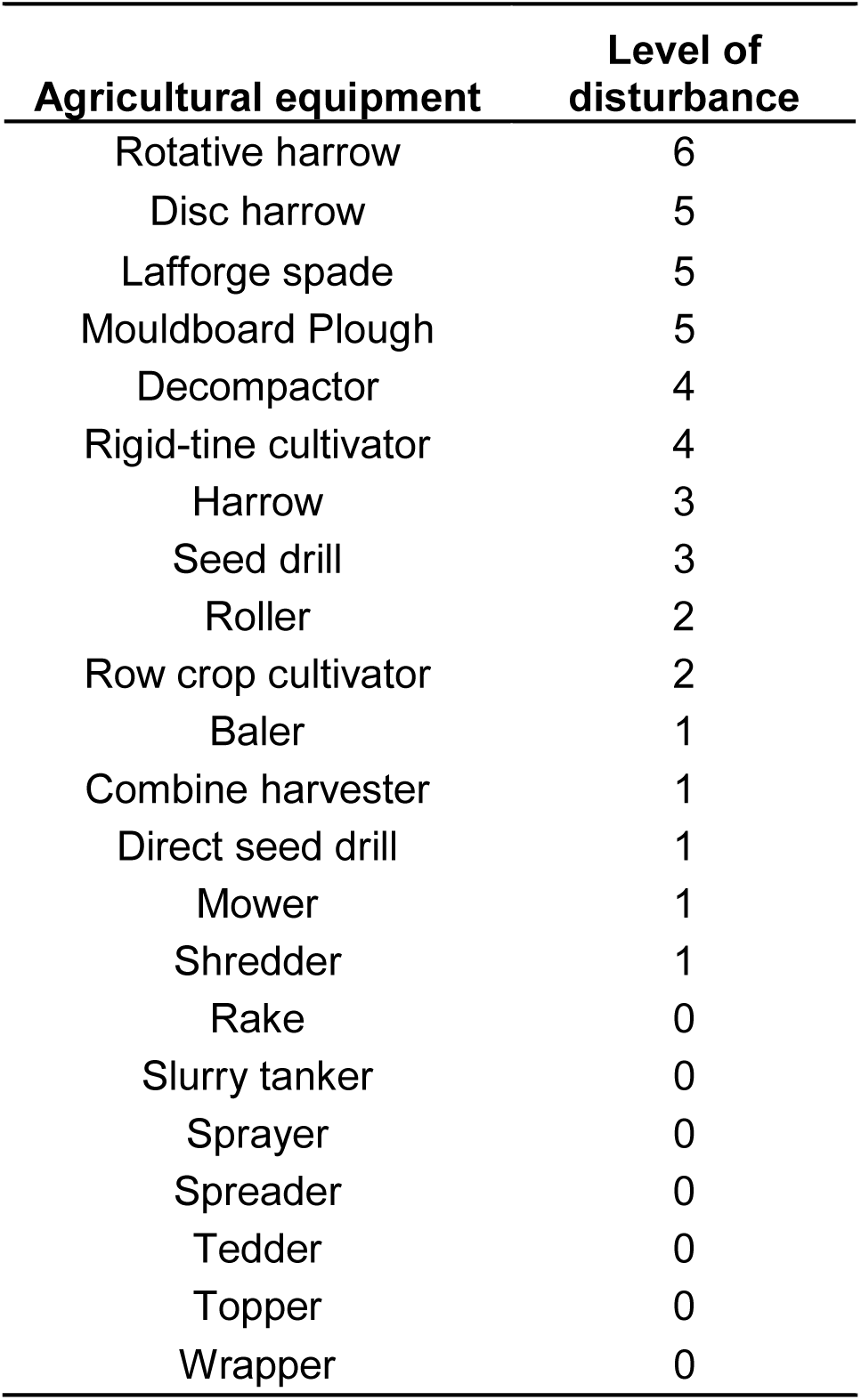
Classification of the agricultural equipment according to their expected effect on the physical habitat stability. The more animated and deeper, the more disturbing.

| **Agricultural equipment** | **Level of disturbance** |
| --- | --- |
| Rotative harrow | 6 |
| Disc harrow | 5 |
| Lafforge spade | 5 |
| Mouldboard Plough | 5 |
| Decompactor | 4 |
| Rigid-tine cultivator | 4 |
| Harrow | 3 |
| Seed drill | 3 |
| Roller | 2 |
| Row crop cultivator | 2 |
| Baler | 1 |
| Combine harvester | 1 |
| Direct seed drill | 1 |
| Mower | 1 |
| Shredder | 1 |
| Rake | 0 |
| Slurry tanker | 0 |
| Sprayer | 0 |
| Spreader | 0 |
| Tedder | 0 |
| Topper | 0 |
| Wrapper | 0 |

**Table S3.**
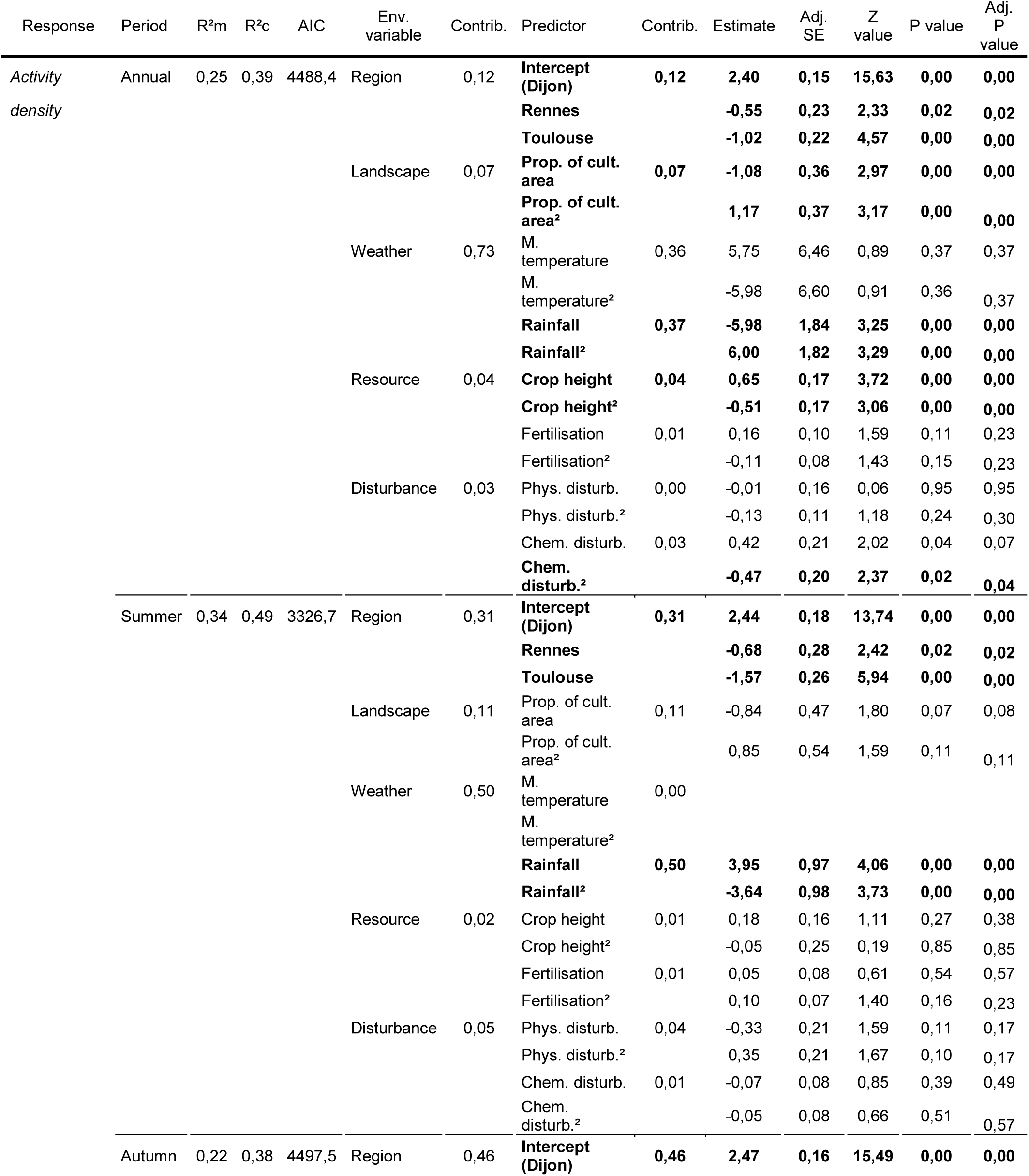
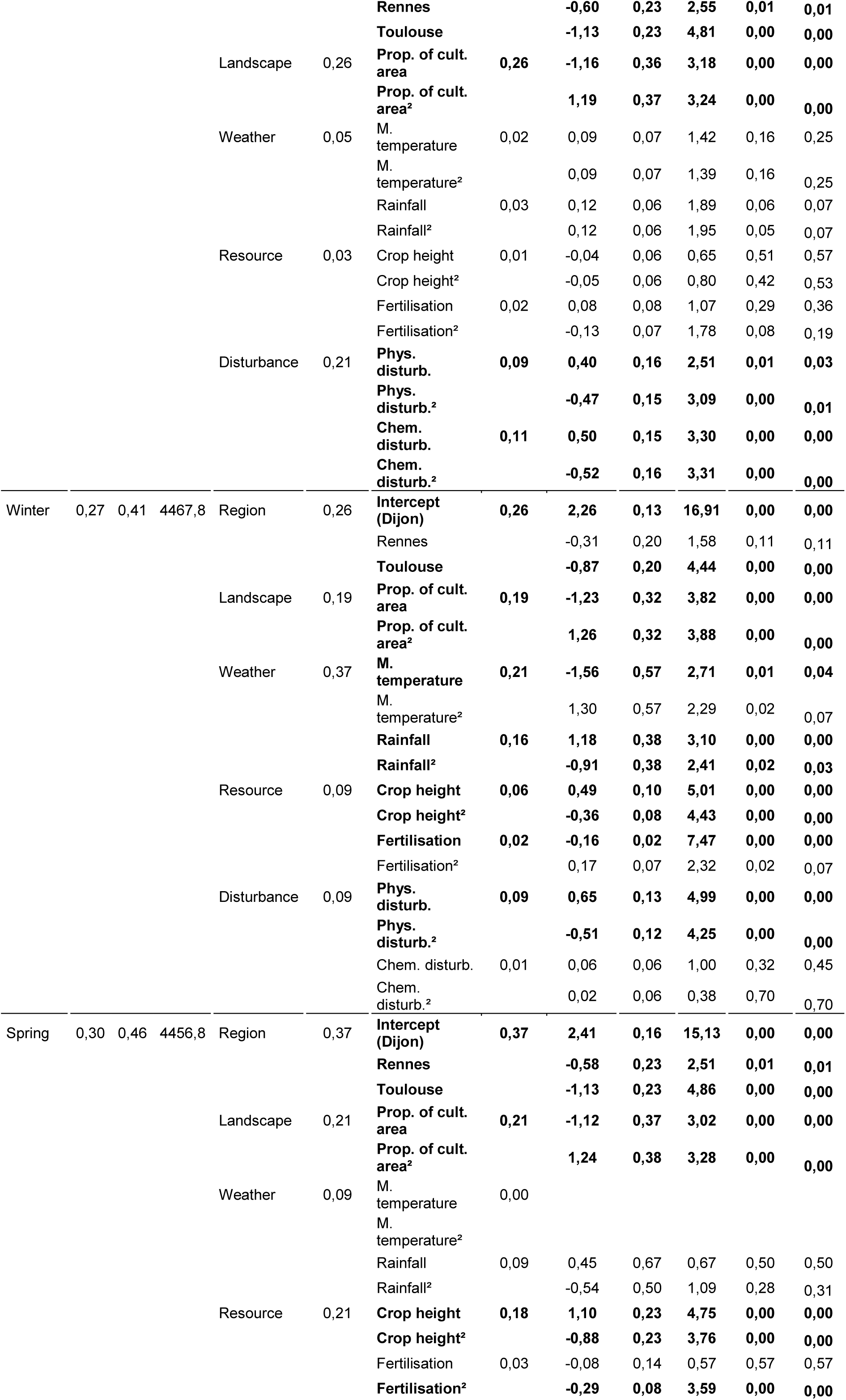
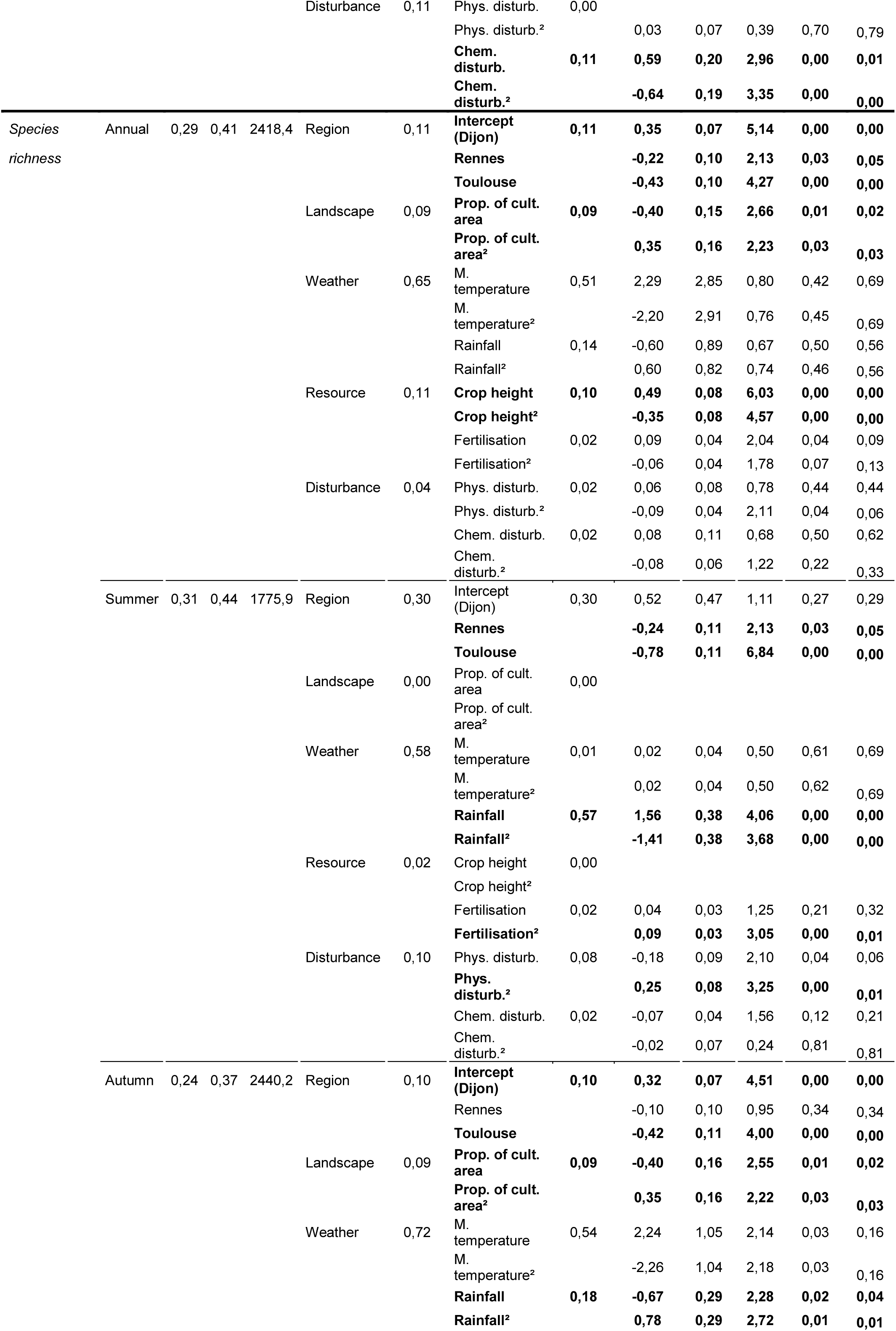
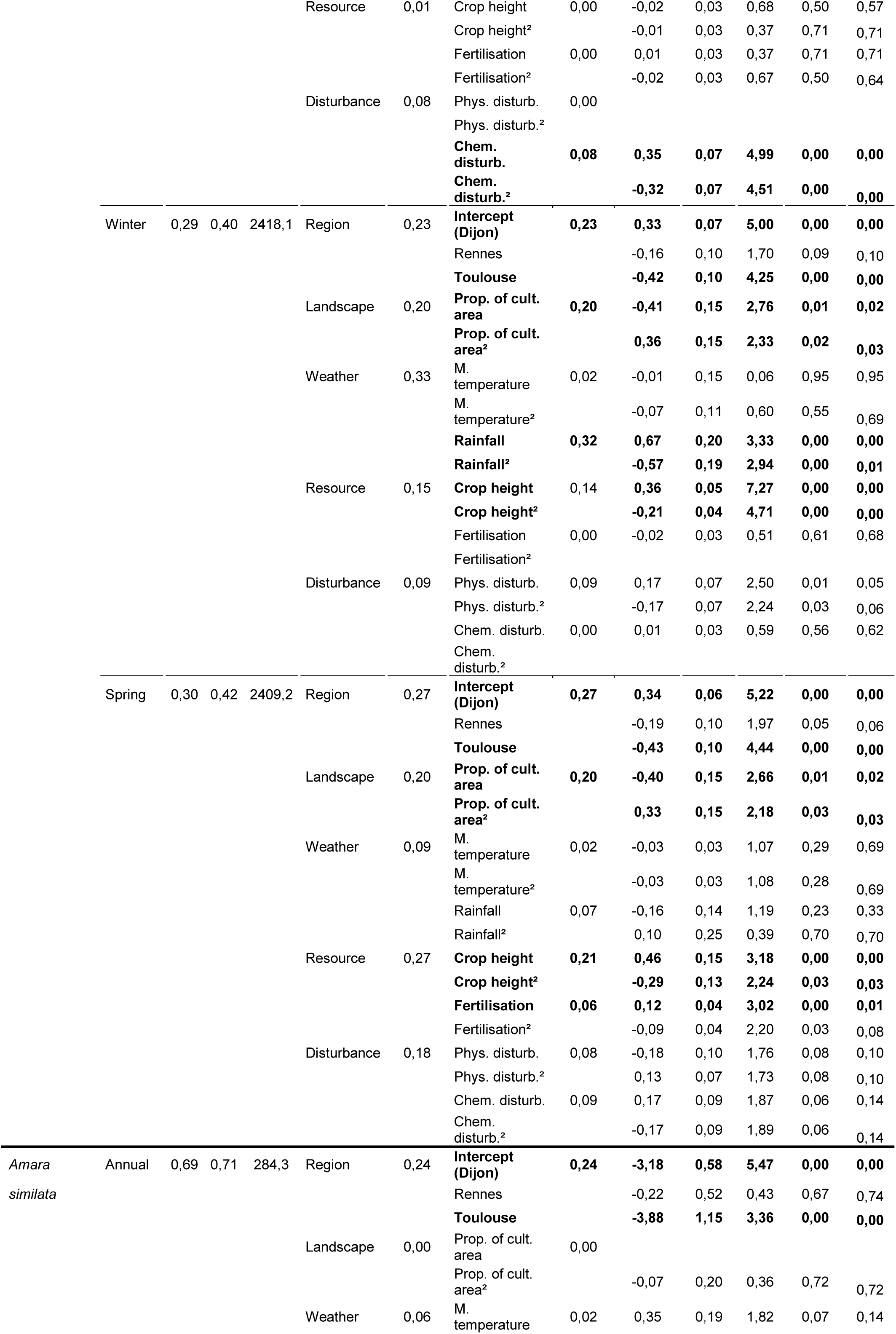
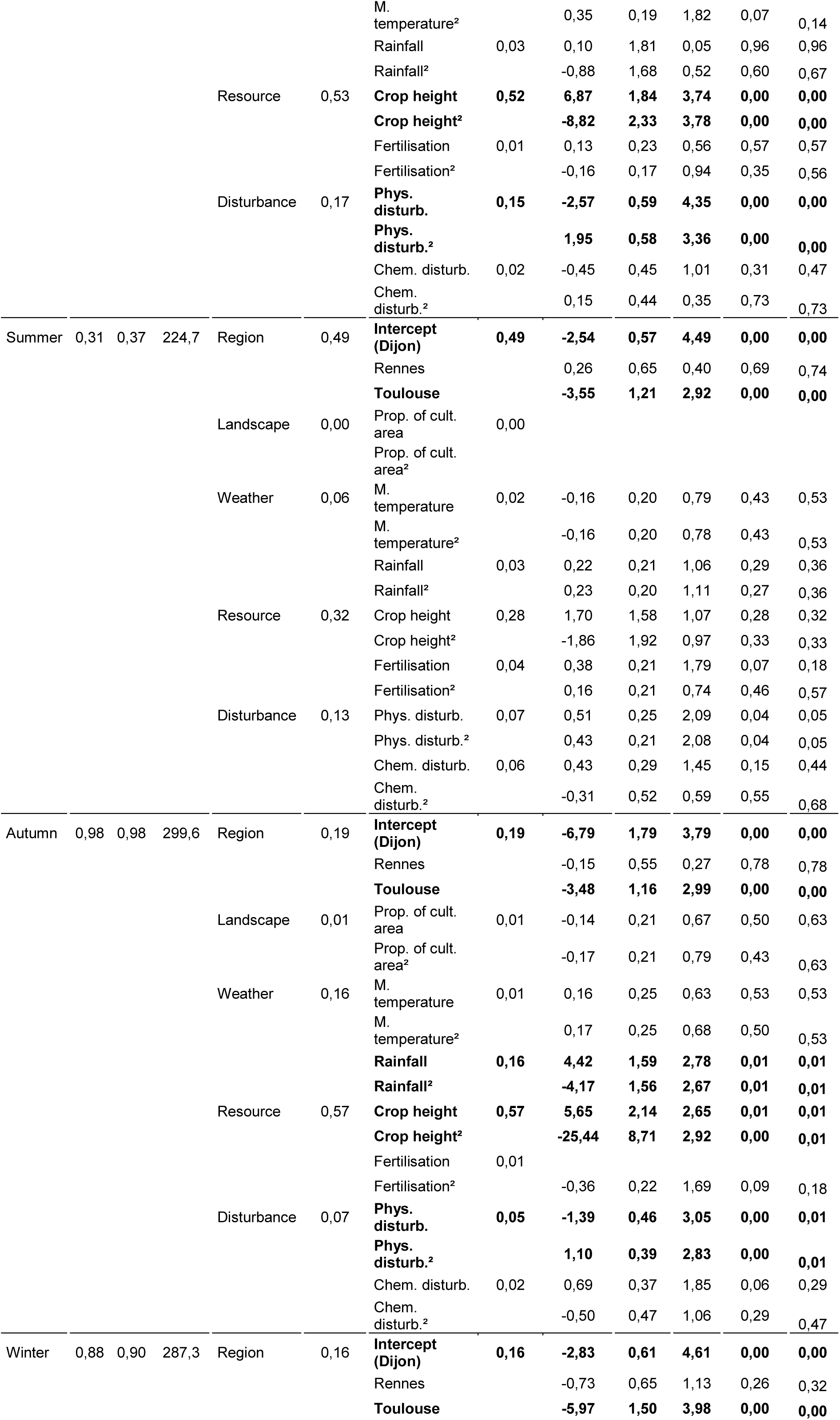
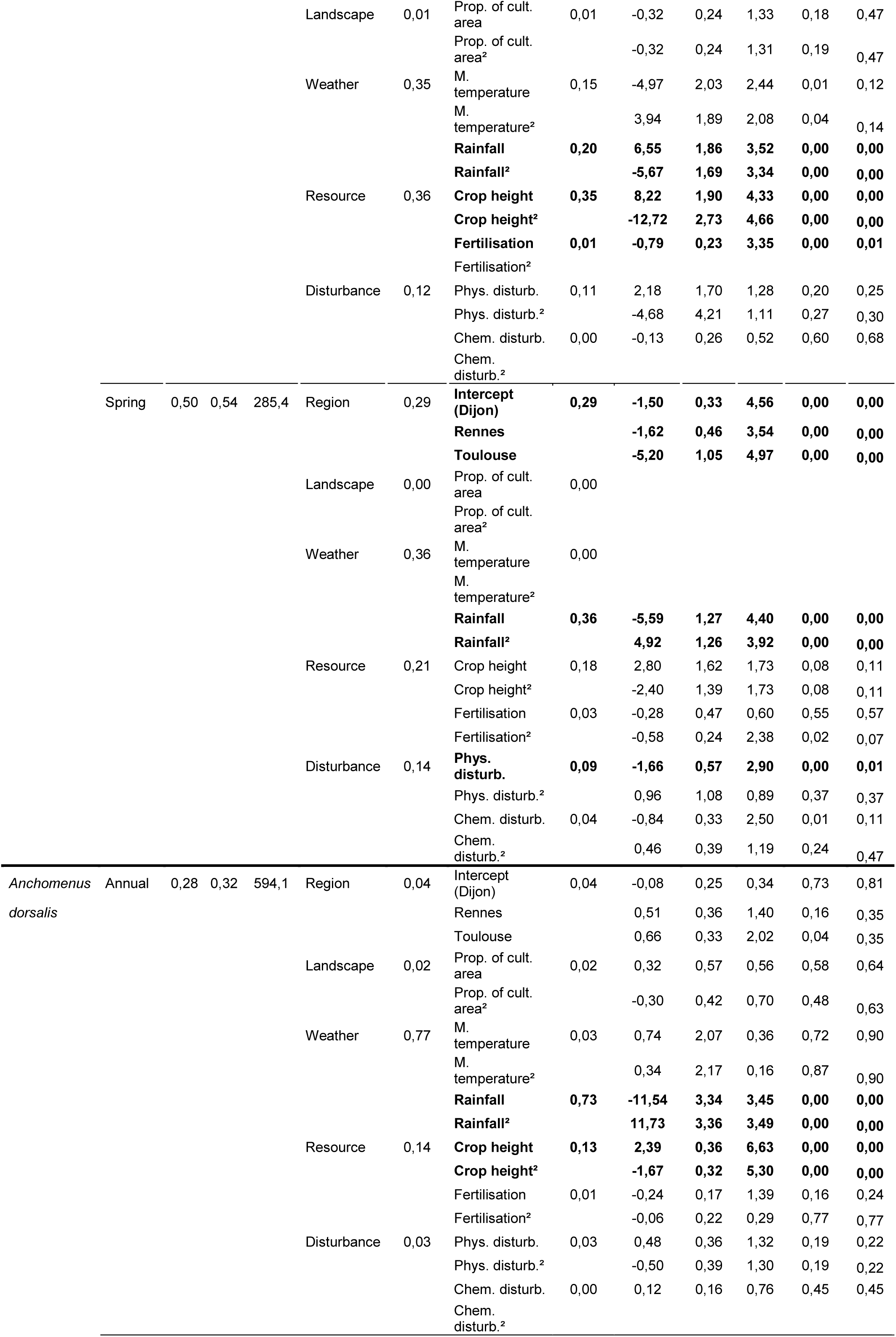
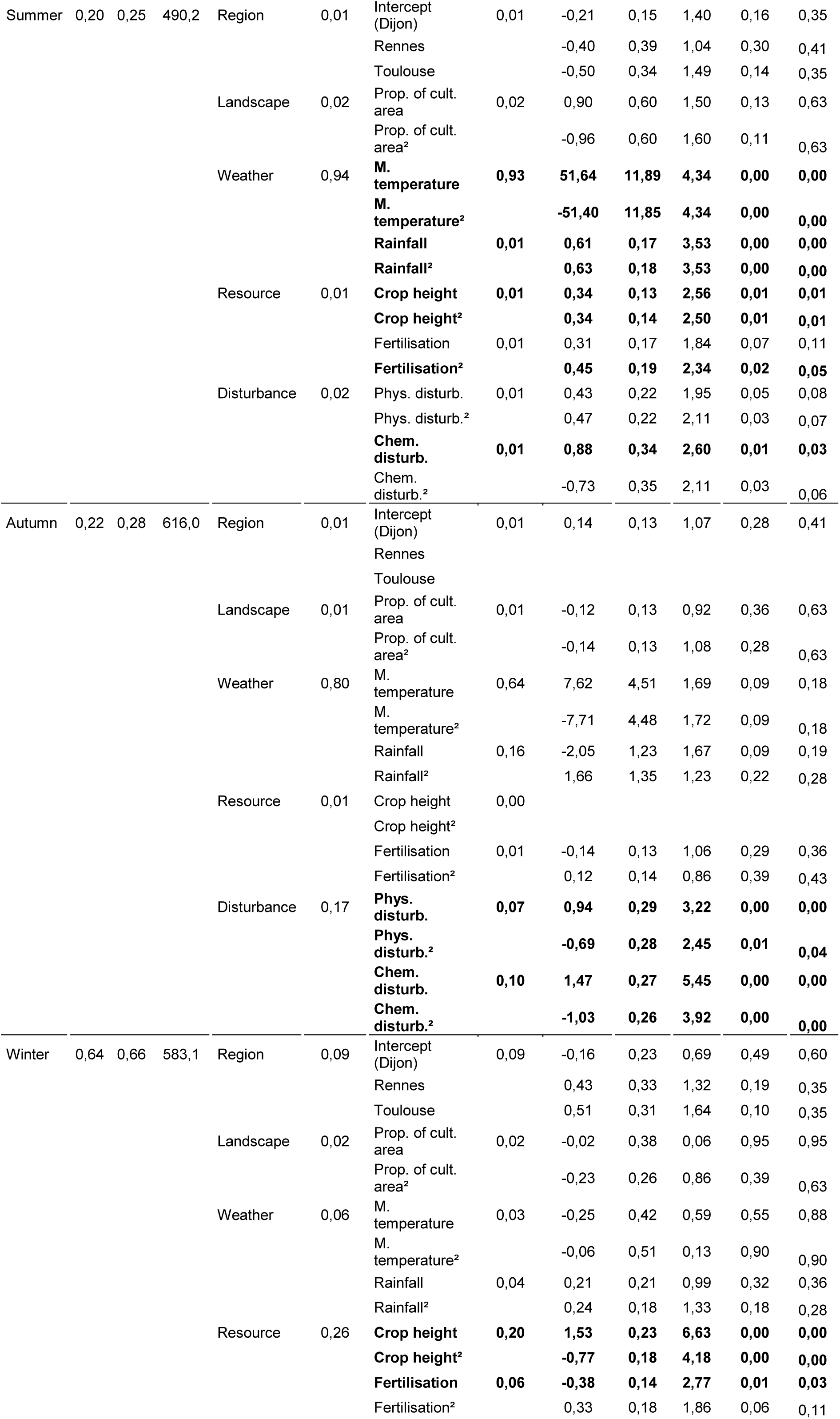
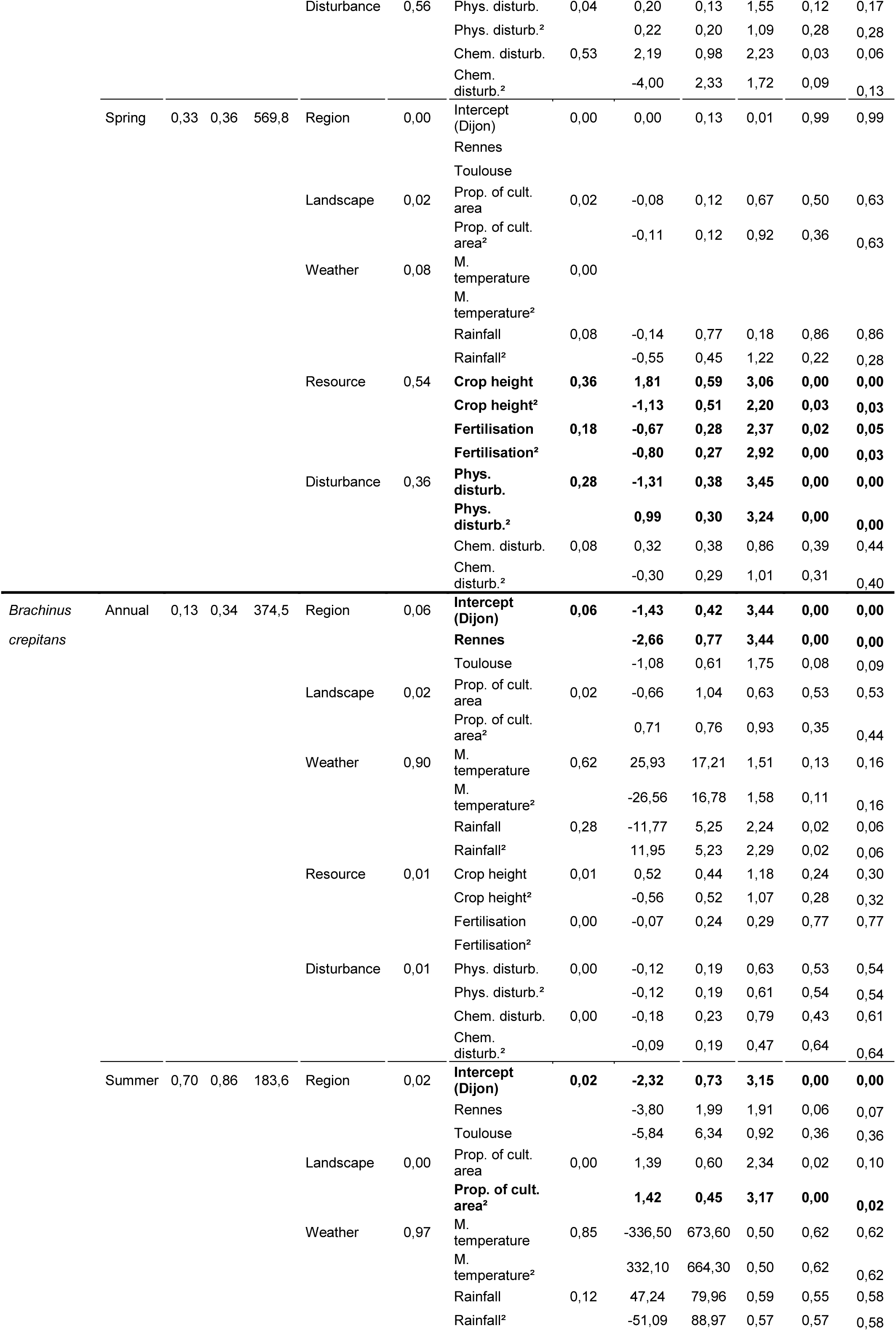
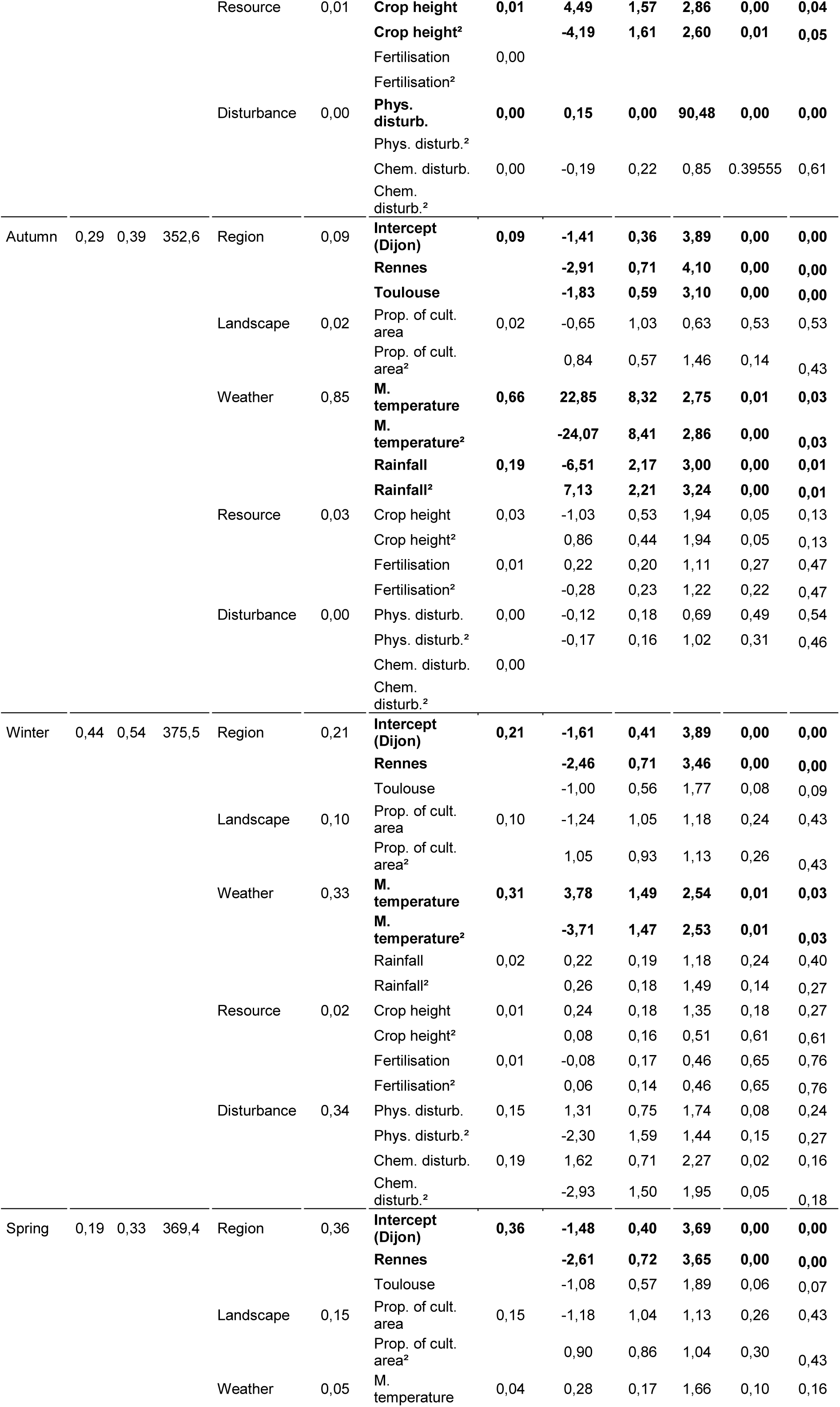
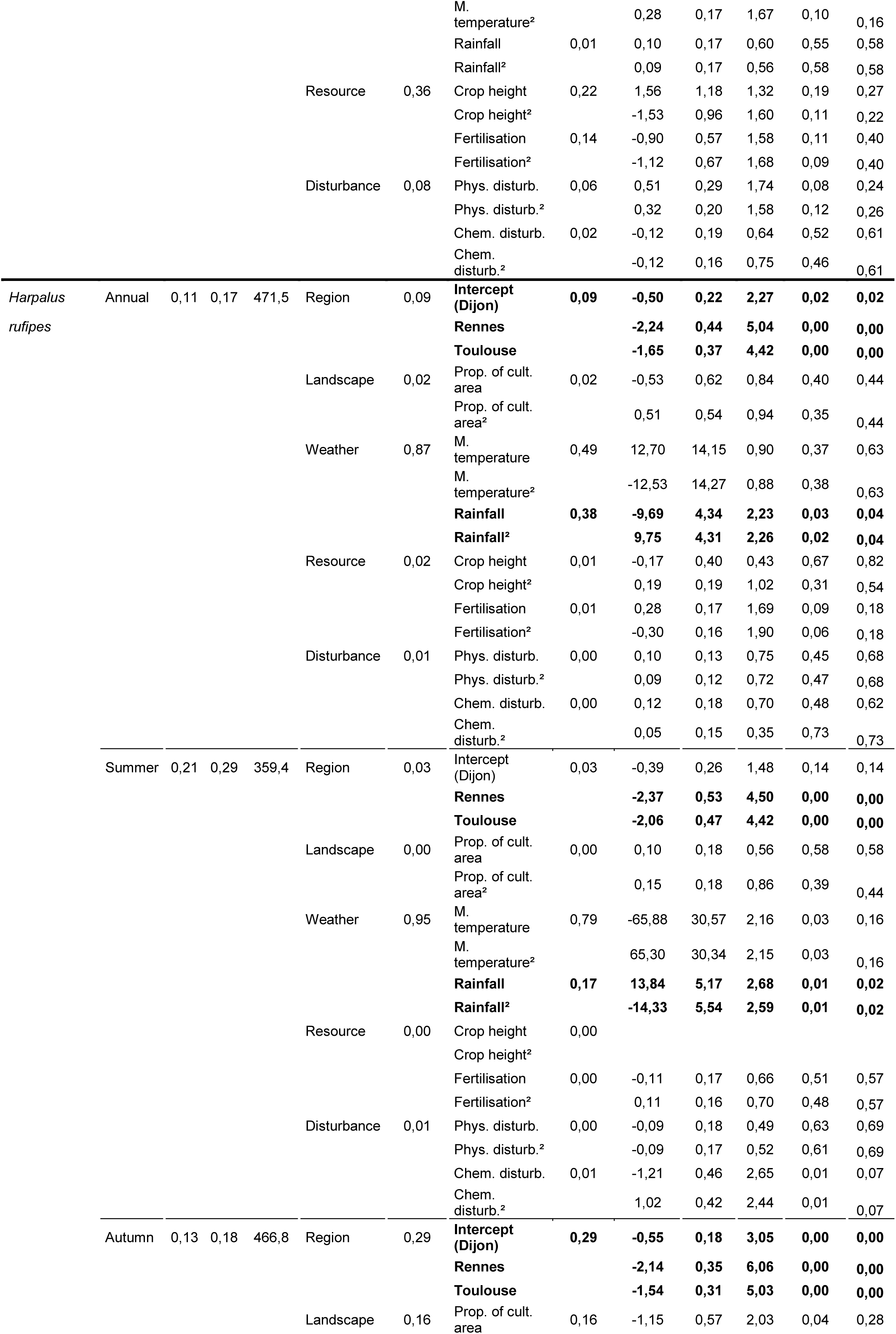
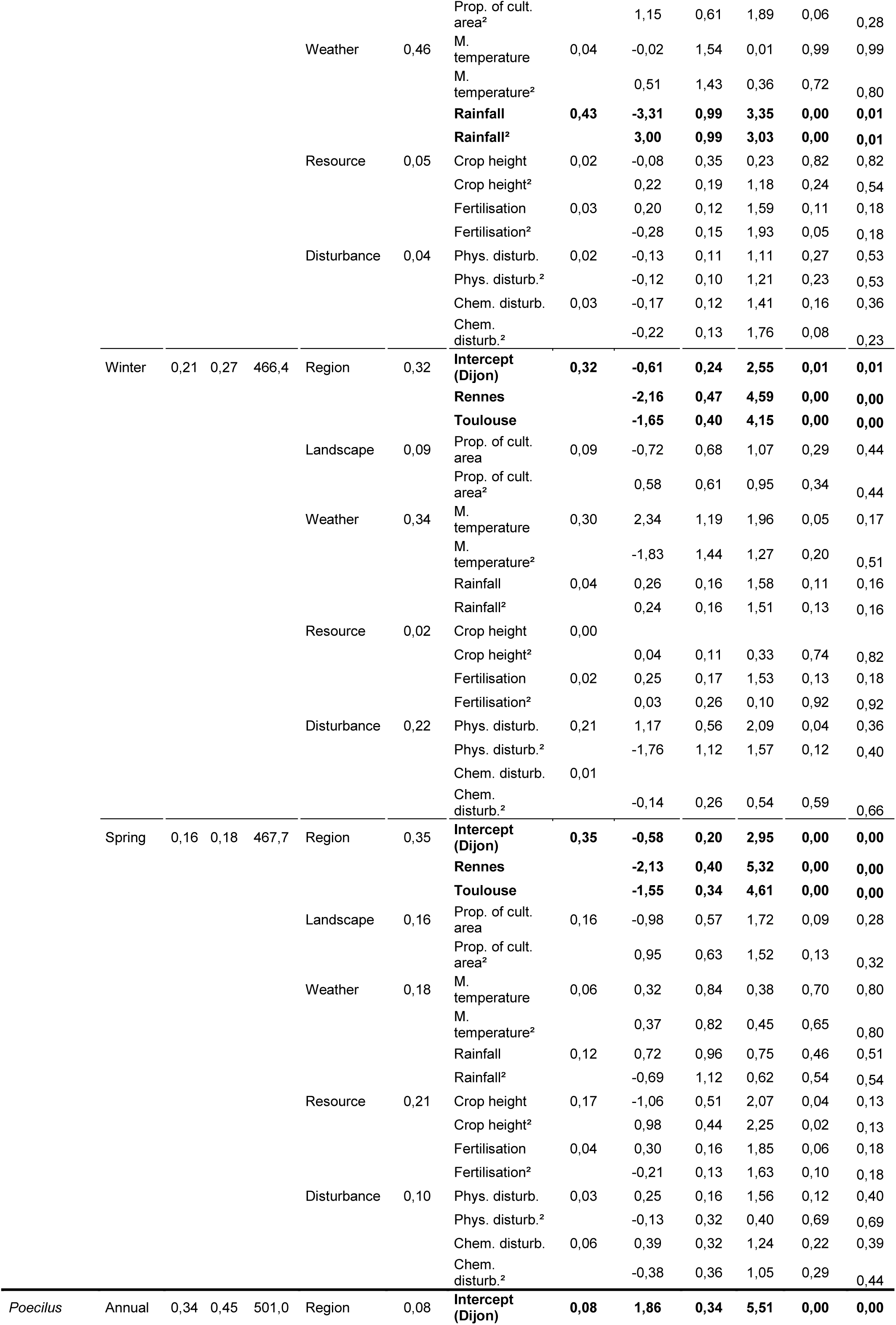
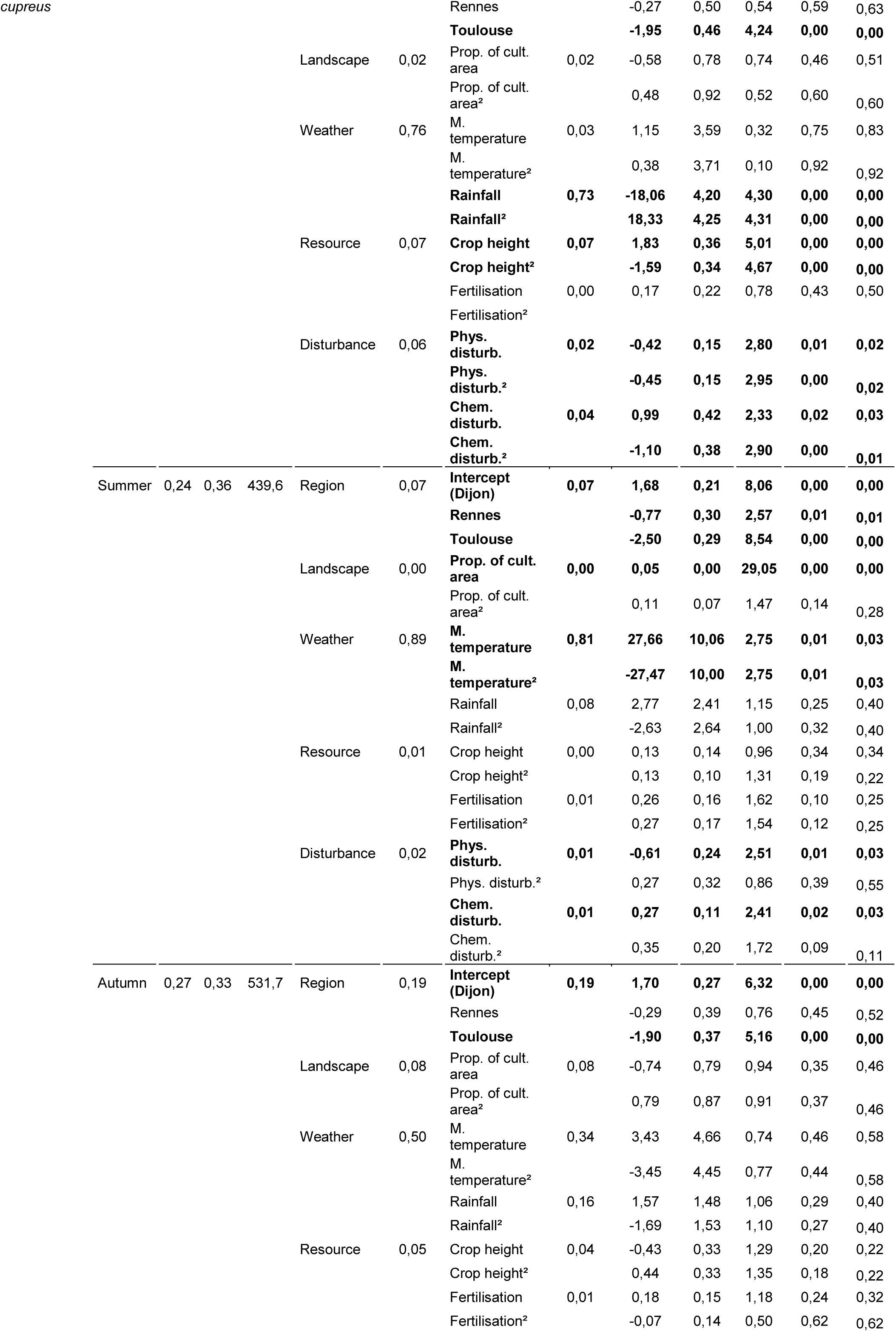
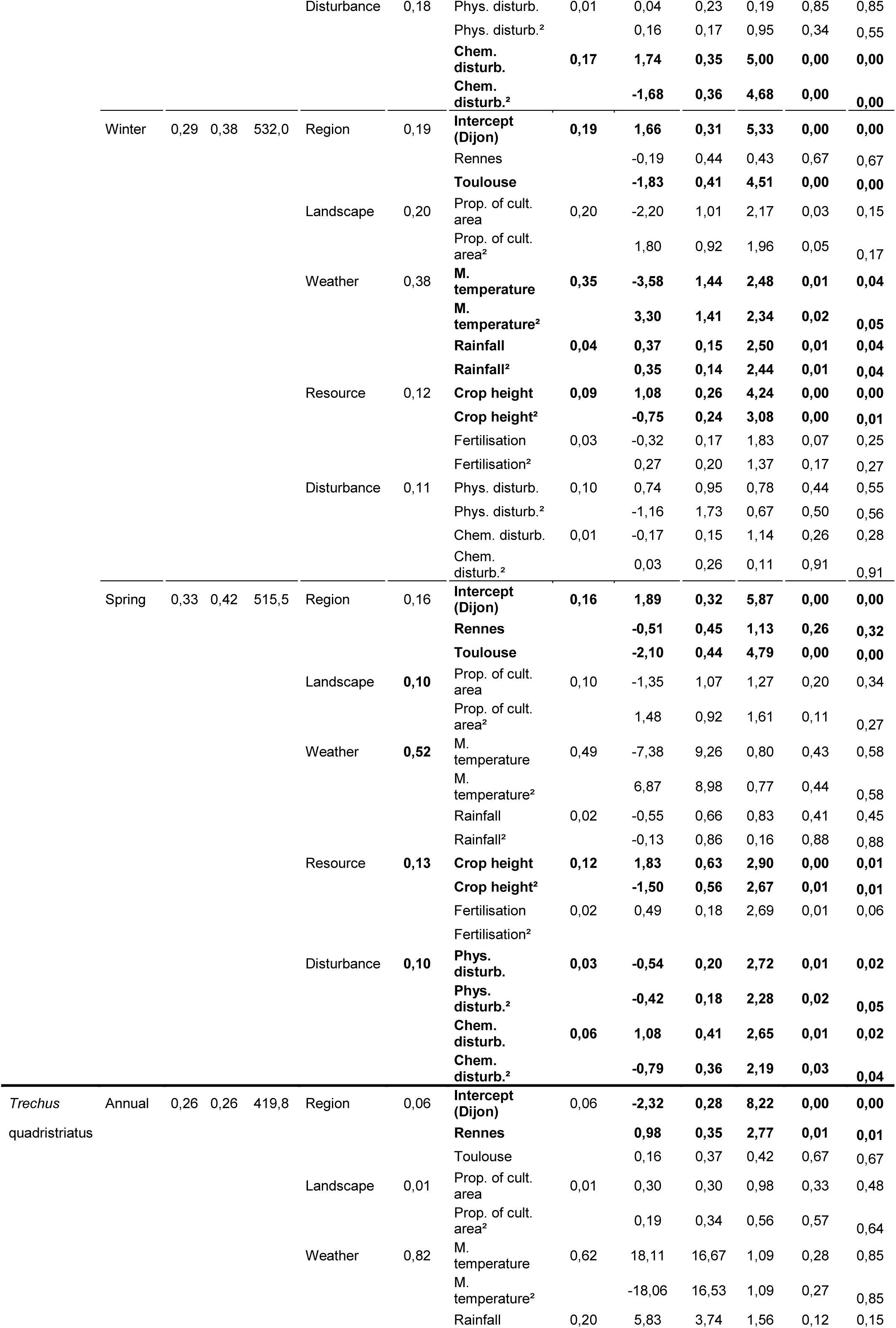
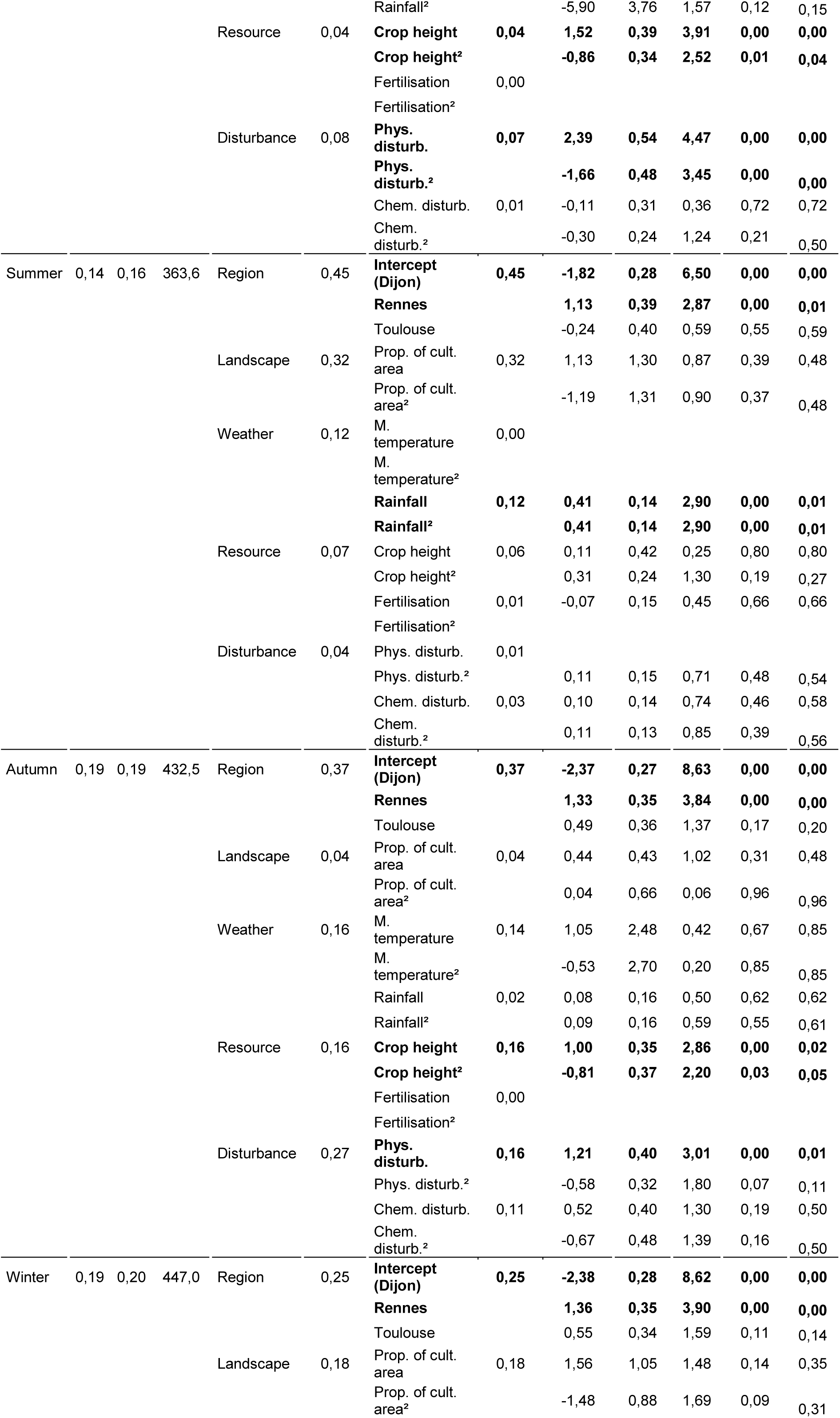
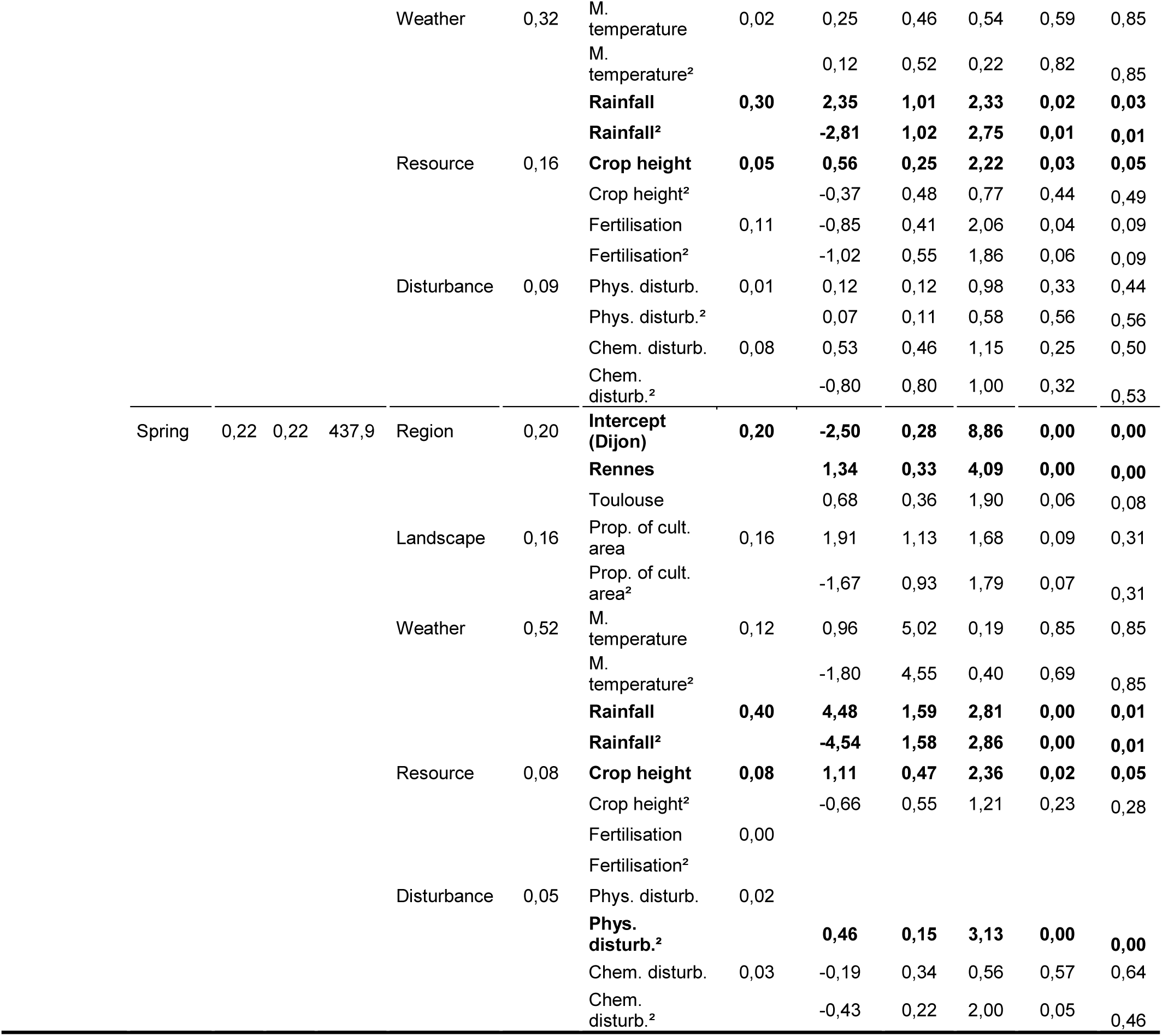
Model selection table for all the response variables and periods. Summer models were based on 390 observations and annual, autumn, winter, spring models on 510 observations. R^2^m are R^2^c were marginal and conditional R^2^ calculated with the delta method from the ‘MuMIn’ package (Bartoń, 2020). ‘Contrib. Feature’ was the contribution of the feature to the explained variance of the model and ‘Contrib pred.’ was the contribution of each predictor to the explained variance of the model. ‘Estim.’, ‘StdErr.’, ‘Z val.’, ‘Pval.’ and ‘Adj. P val’ were estimates, standard error, Z-value, P-values and adjusted P-values associated to each predictor. Adj. P-values were estimated with the « false discovery rate » method (Benjamini et al., 2001). Predictors were considered as significant were their adjusted p-values was below 0.05, they were written in bold below. Abbreviation of predictors: ‘Prop. of cult. area’ was ‘proportion of cultivated area’ ; ‘Mean temp.’ was ‘Mean temperature’ ; ‘Phys. disturb’ was ‘Physical disturbance’ and ‘Chem. disturb.’ was ‘Chemical disturbance’. When Estimates, Standard error, Z values and P values were missing for a predictor, it means that itw as not retained by the model selection procedure.

**Figure S1.**
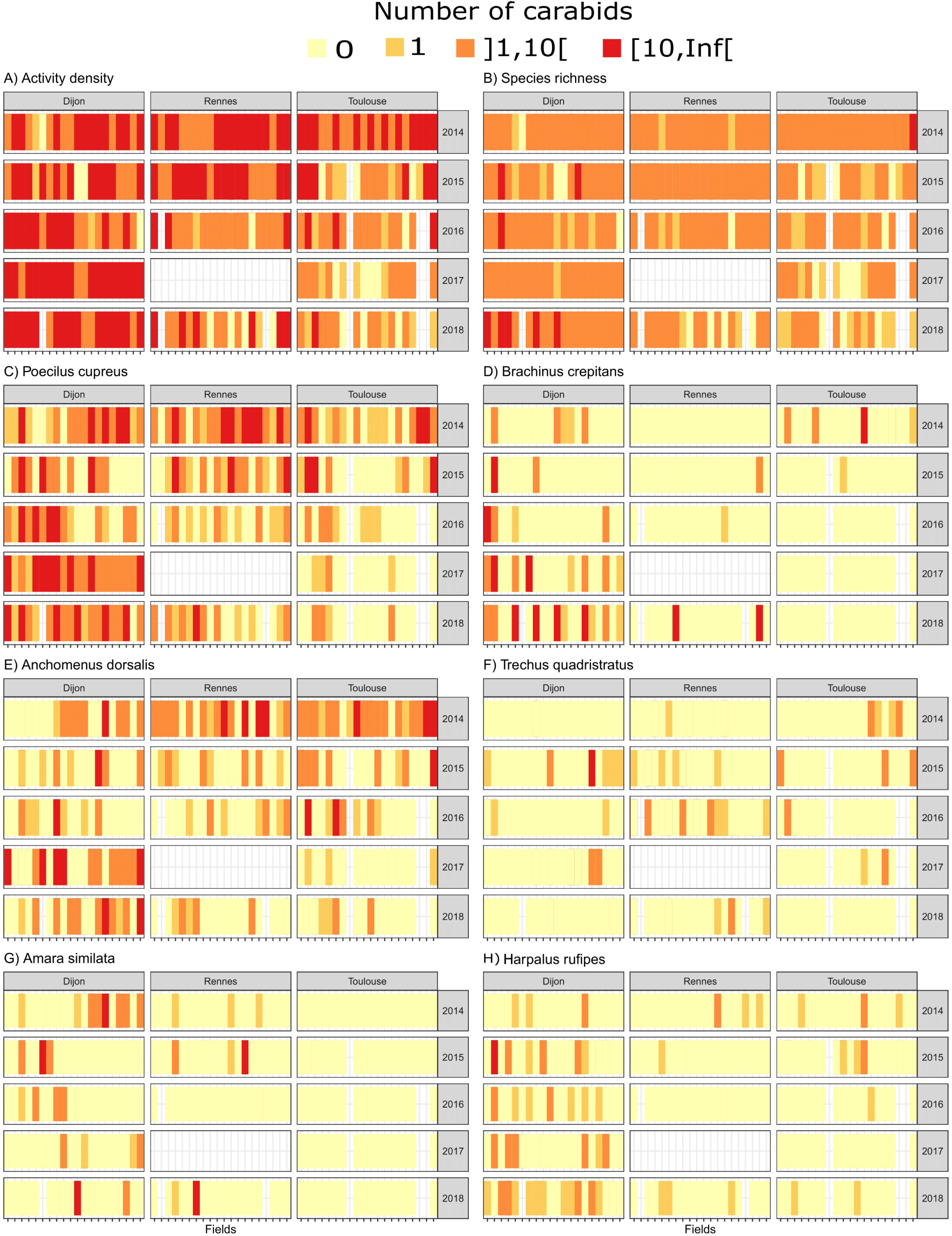
Values of the five carabid response variables region, year and field. For each year and region, values for the twenty fields are presented into four classes represented by distinct colours. Annual carabid data per field were derived from 16 pitfall traps (8 traps at two sampling dates). In some instances, carabids could not be sampled in some fields (represented in white).

**Figure S2.**
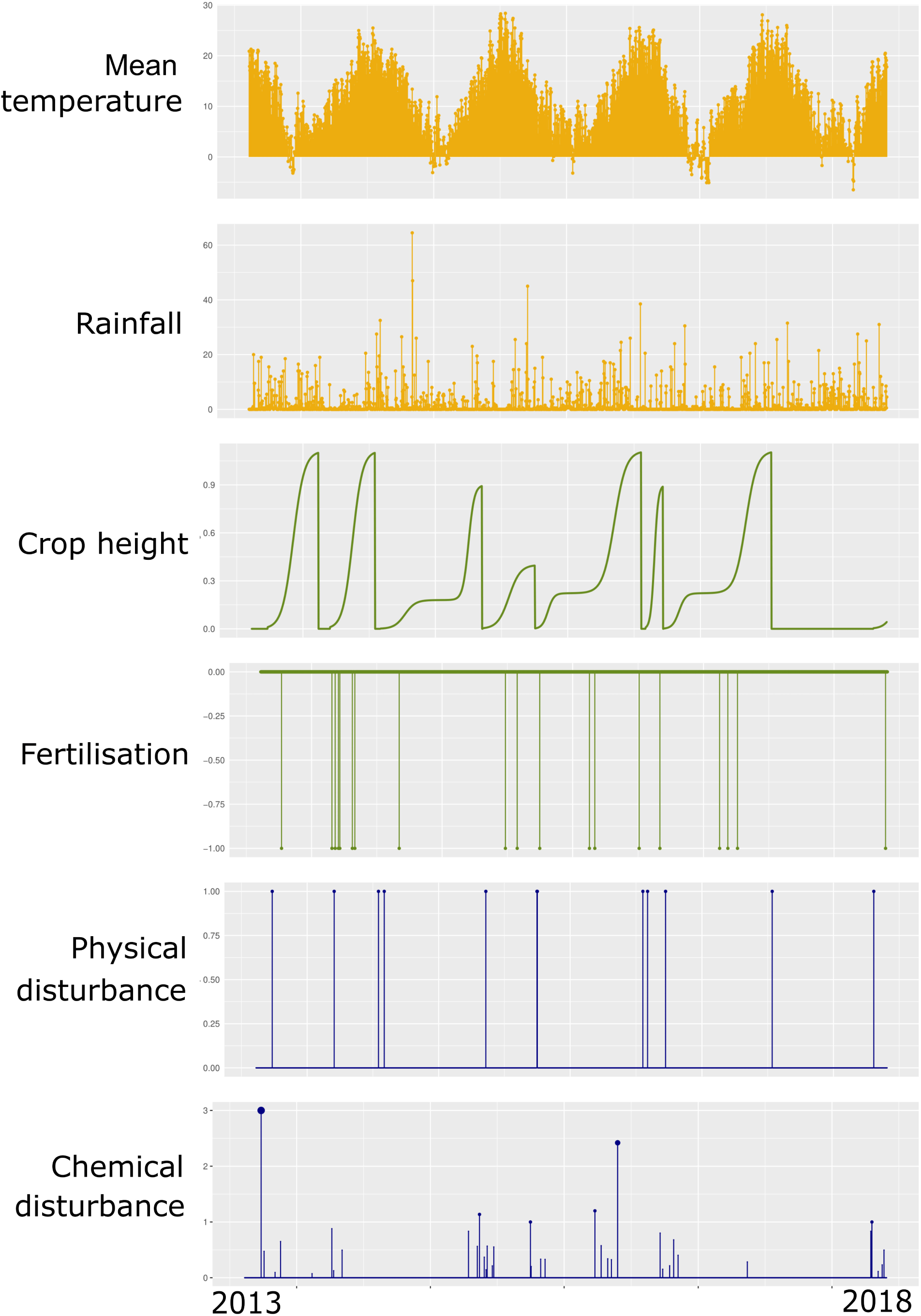
A representation of the conversion of meteorological data and technical routes data into ecological gradients over the 5-years period for a field. This example represents a field conducted under conservation agriculture.

**Figure S3.**
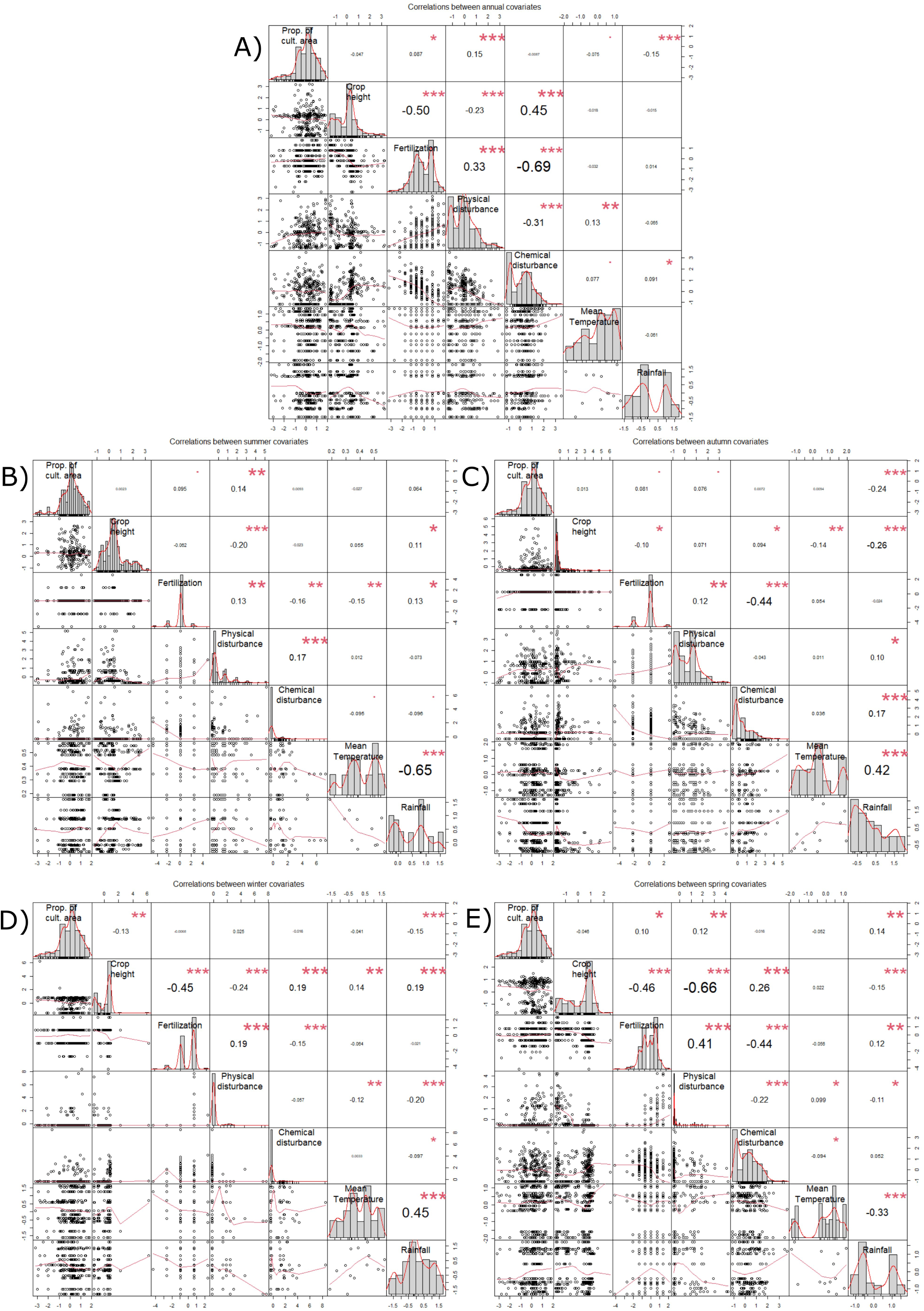
Correlation matrix between all the explanatory variables for each period (A. Annual, B. Summer, C. Autumn, D. Winter, E. Spring).

**Figure S4.**
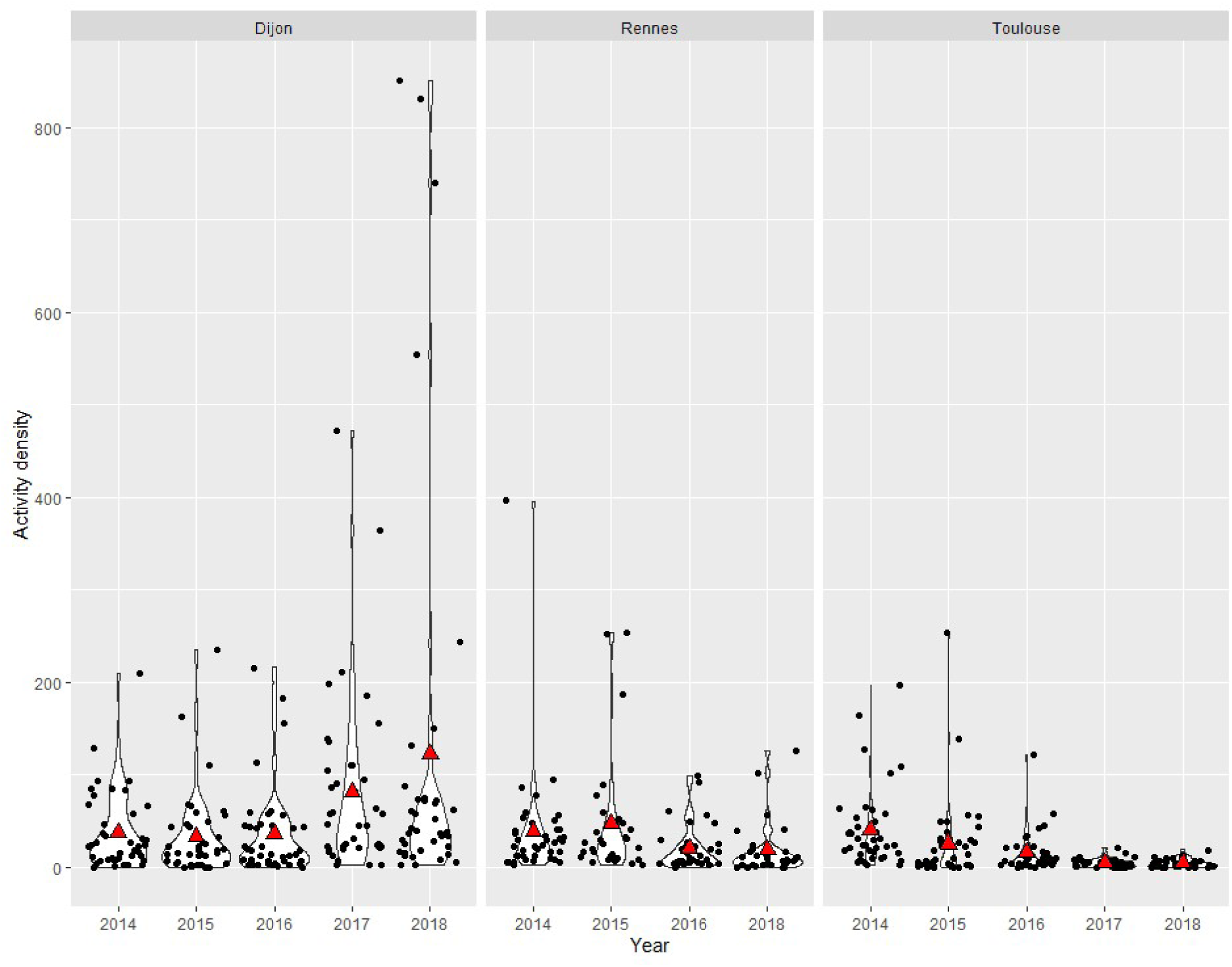
Temporal patterns of the activity density of carabids in the three regions. Each point represents a sampling point for a field. Red triangles represent the mean activity density for the given year and region.

**Figure S5.**
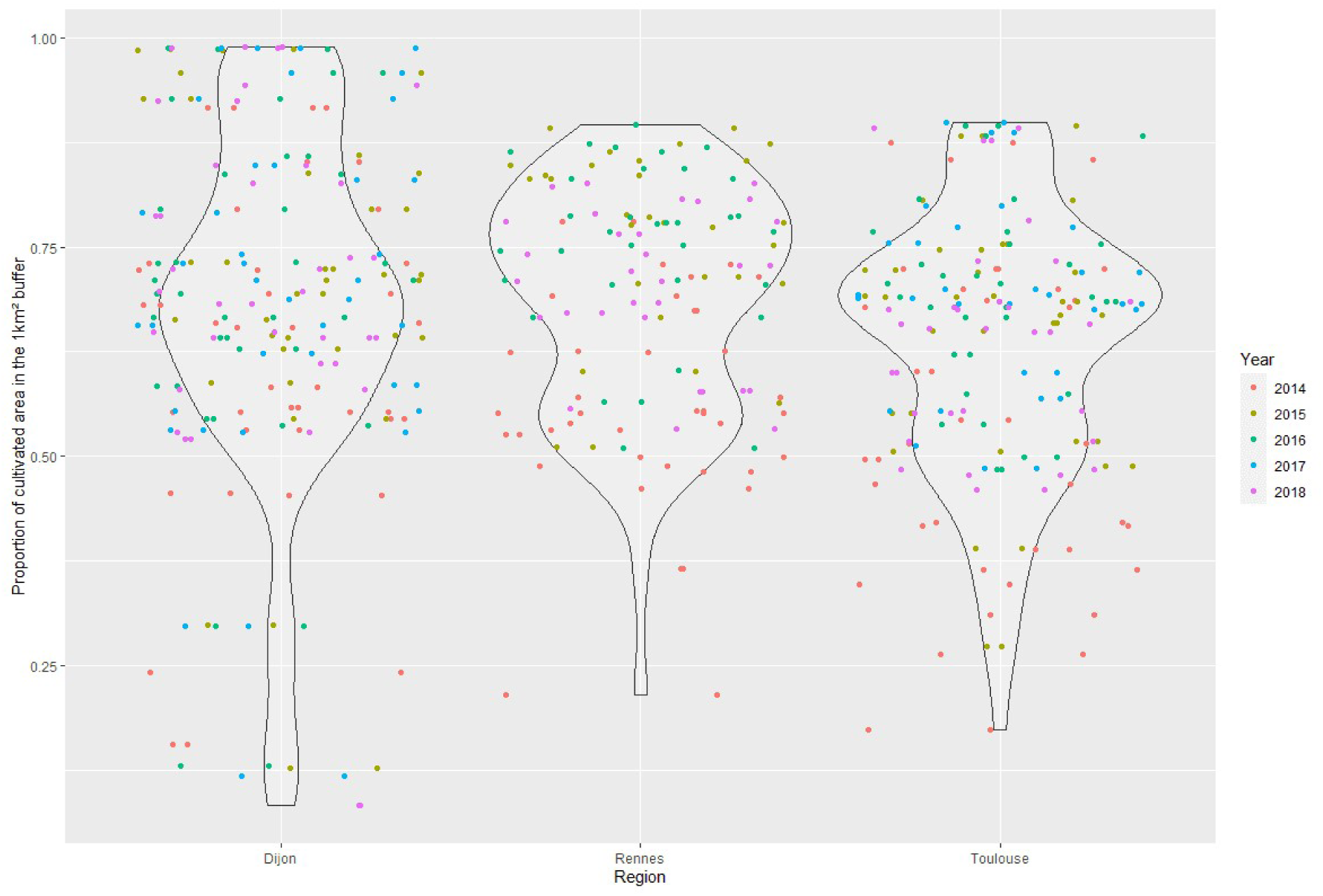
Distribution of the proportion of cultivated area in the 1km^2^ buffer around each focal field for each year.

### Annexe 1 Modelling the growth of different crops

Three types of curves were used to model crop growth. A logistic growth was used for crops for which the growth period did not include the winter period (period defined from the 15th of December to the 15h of February), following the equation:

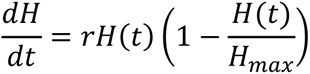

with *r* such as *H* = *H_max_* at *t* = harvest date. For winter crops and some intercrops for which the growth period included the winter period, we used the same logistic equation but with a plateau between the 15th of December and the 15th of February corresponding to a break in the vegetative growth (the crop height at the plateau was arbitrary chosen as 0.2 × *H_max_*). Finally, grazing periods were considered as a linear decrease of crop height from the first day of grazing over 15 days long (estimated from the expertise of agronomists because we did not collect information on grazing duration).

### Annexe 2 How to identify biological thresholds with quadratic effects

The turning point (interpreted as a biological threshold) of each predictor having a quadratic effect is obtained by finding the root of the derivative of the quadratic function. Below, *y(x)* is the response variable with respect to the predictor *x* . We also denote by *a the* intercept, *b* and *c* the parameters of first and second order of the predictor. Furthermore, the magnitude of c reveals the magnitude of the quadratic effect.

More precisely, we have

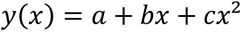

#### The derivative with respect to x writes

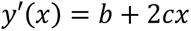

#### From where we obtain the root of the derivative (called ‘turning point’)

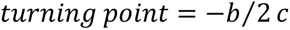

**Figure Annexe 2.**
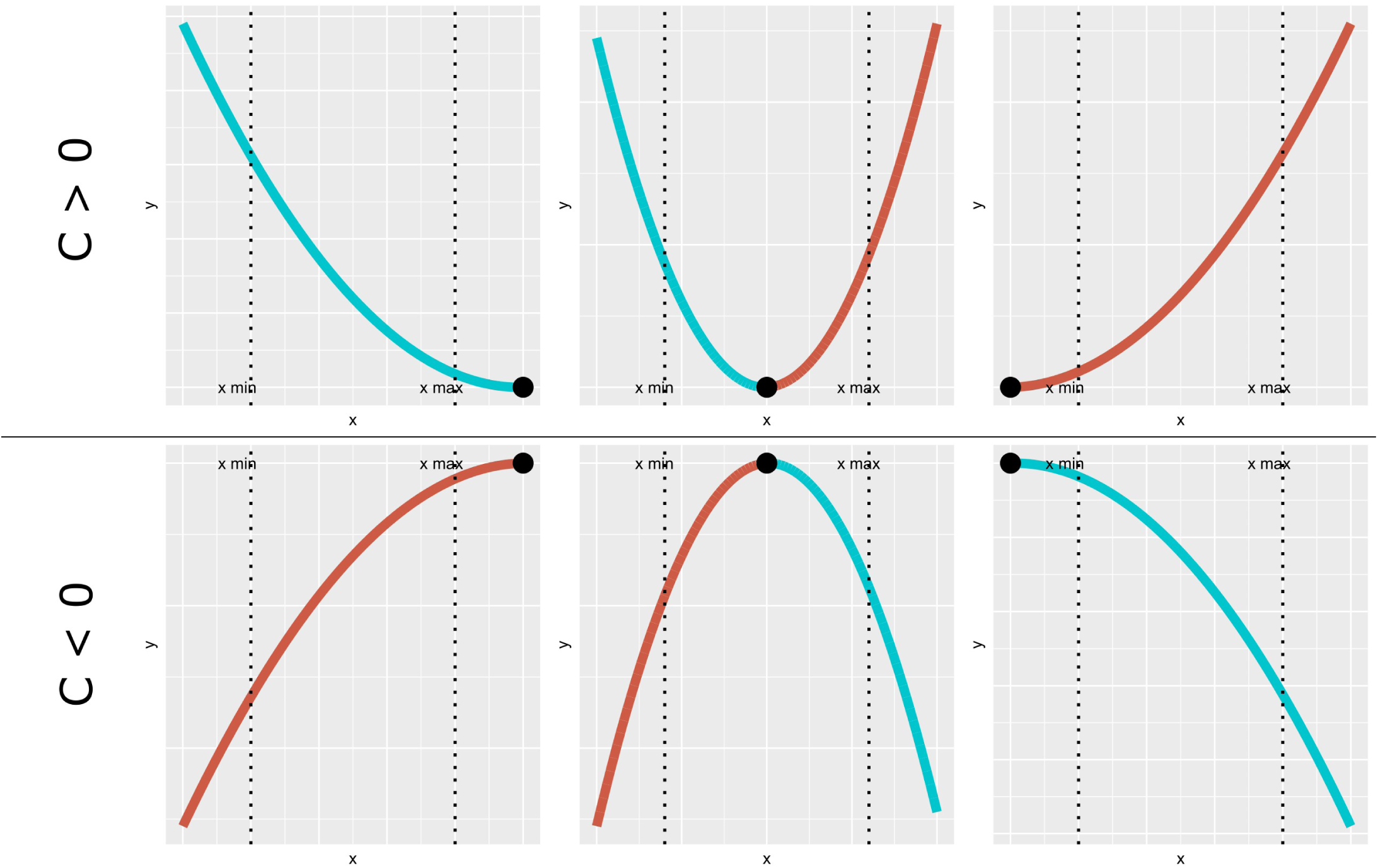
Determination of the relationship between x and y=a+bx+cx^2^ depending on the range of x values and the turning point (-b/2c). The inflection point is identified with a black point in each panel. Blue curves represent a negative effect of x on y while red curves represent a positive effect.

### Annexe 3 Variations of resource and disturbance gradients over seasons

As expected, crop height is minimal in autumn and progressively increased until summer (Table and figure of this annexe). Fertilisers were rarely spread in summer and autumn and the number of spreading increased in winter and spring. The level of physical disturbances decreased from autumn, to summer, spring and winter. Finally, the level of chemical disturbances was very low in summer and winter and increased in autumn and spring.

**Table Annexe 3.**
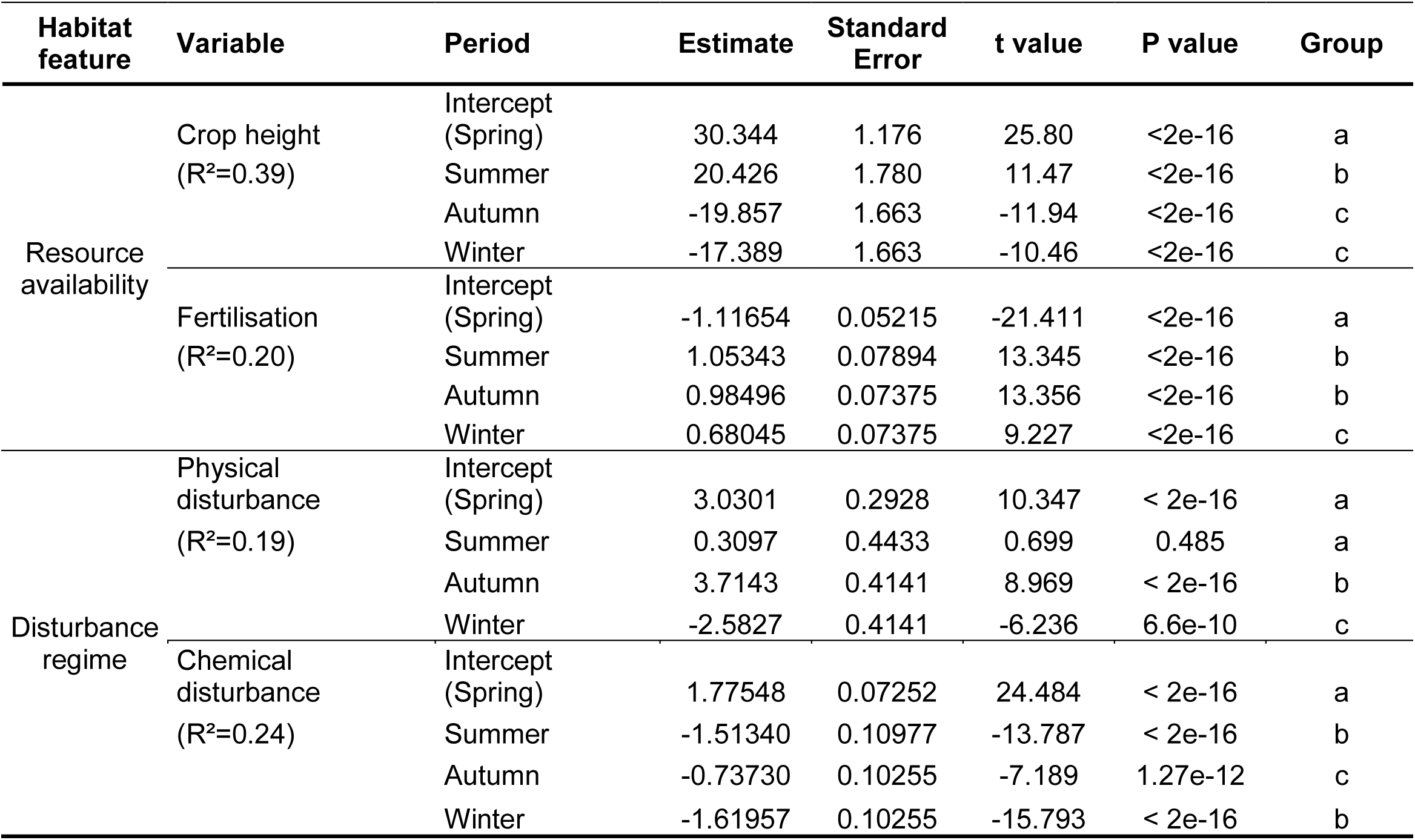
Effect of the season on explanatory variable of the mixed effect models. Explanatory variables varied according to seasons. Estimate, Standard error, t value and p-values were obtained with an ANOVA. Group were defined using a Tukey Test.

| Habitat feature | Variable | Period | Estimate | Standard Error | t value | P value | Group |
| --- | --- | --- | --- | --- | --- | --- | --- |
| Resource availability | Crop height<br>(R<sup>2</sup>=0.39) | Intercept (Spring) | 30.344 | 1.176 | 25.80 | <2e-16 | a |
|  |  | Summer | 20.426 | 1.780 | 11.47 | <2e-16 | b |
|  |  | Autumn | -19.857 | 1.663 | -11.94 | <2e-16 | c |
|  |  | Winter | -17.389 | 1.663 | -10.46 | <2e-16 | c |
|  | Fertilisation<br>(R<sup>2</sup>=0.20) | Intercept (Spring) | -1.11654 | 0.05215 | -21.411 | <2e-16 | a |
|  |  | Summer | 1.05343 | 0.07894 | 13.345 | <2e-16 | b |
|  |  | Autumn | 0.98496 | 0.07375 | 13.356 | <2e-16 | b |
|  |  | Winter | 0.68045 | 0.07375 | 9.227 | <2e-16 | c |
| Disturbance regime | Physical disturbance<br>(R<sup>2</sup>=0.19) | Intercept (Spring) | 3.0301 | 0.2928 | 10.347 | < 2e-16 | a |
|  |  | Summer | 0.3097 | 0.4433 | 0.699 | 0.485 | a |
|  |  | Autumn | 3.7143 | 0.4141 | 8.969 | < 2e-16 | b |
|  |  | Winter | -2.5827 | 0.4141 | -6.236 | 6.6e-10 | c |
|  | Chemical disturbance<br>(R<sup>2</sup>=0.24) | Intercept (Spring) | 1.77548 | 0.07252 | 24.484 | < 2e-16 | a |
|  |  | Summer | -1.51340 | 0.10977 | -13.787 | < 2e-16 | b |
|  |  | Autumn | -0.73730 | 0.10255 | -7.189 | 1.27e-12 | c |
|  |  | Winter | -1.61957 | 0.10255 | -15.793 | < 2e-16 | b |

**Figure Annexe 3.**
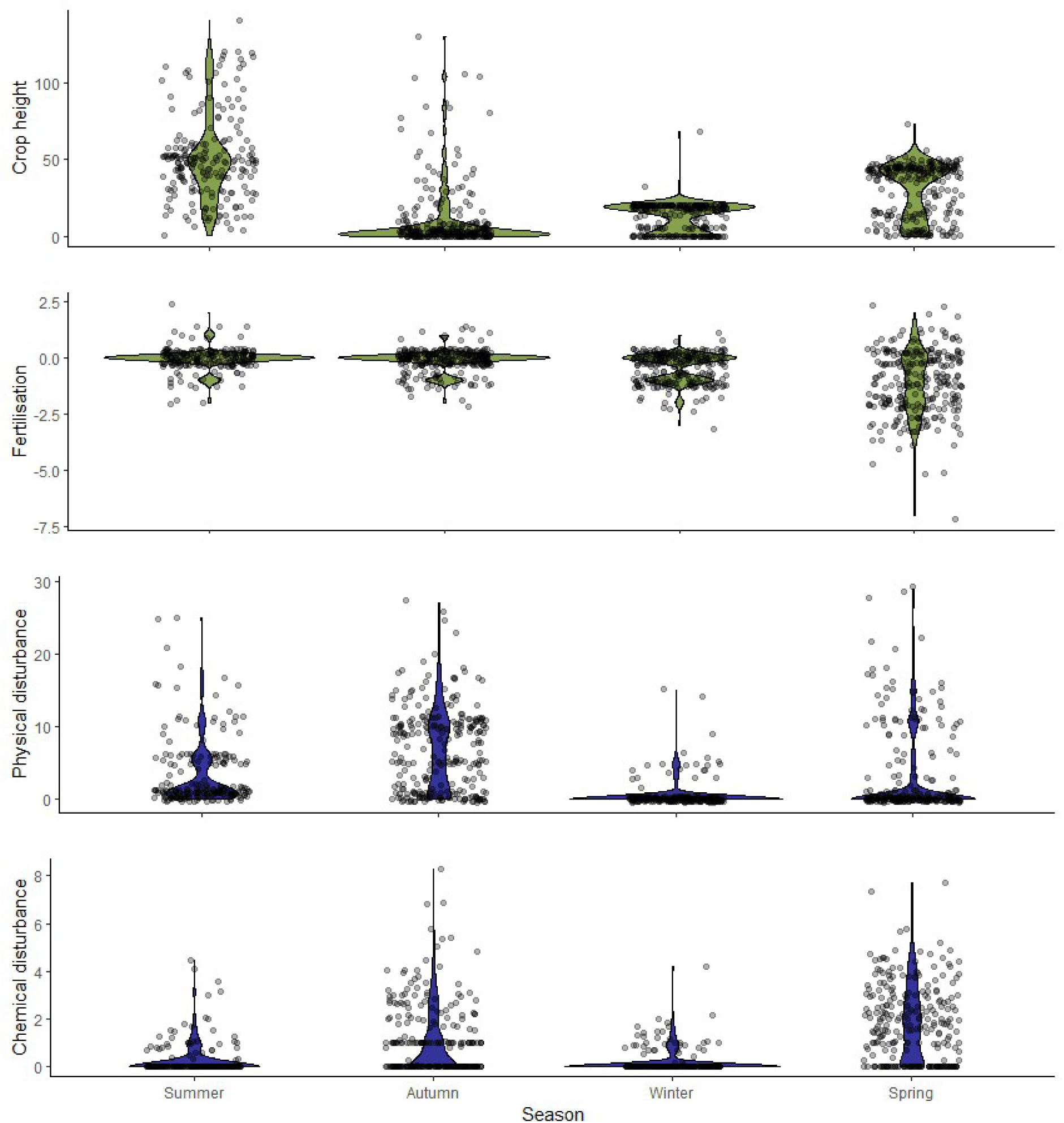
**Seasonal variations of resource availability** (Crop height and Fertilisation) **and disturbance intensity** (Physical disturbance and Chemical disturbance).

### Annexe 4 Effects of the environmental context on carabid distributions

#### Supplementary results

Meteorological conditions were the most influencing variable (Figure 3 of the main text). On average, it explained about 46% of variations in carabid distributions among all the set of best models with a minimum of 20% for *A. similata* and a maximum of 62% for *B. crepitans* (Figure of this annexe and Table S3). Meteorological conditions influenced the most carabid distributions in annual models (70%). Mean temperature contributed more to explain carabid distributions than rainfall (27% vs 20% respectively) but rainfall was significant in a higher number of models than mean temperature (20 vs 6 models). First, the main effect of mean temperature was positive and detected on *P. cupreus* and *H. rufipes* in summer models and *H. rufipes* in autumn and winter models. In addition, winter mean temperature had a negative effect on *P.cupreus* and a hump shaped effect on activity density. Second, rainfall had contrasting effects on carabid depending on periods and response variables. For annual models, rainfall was negatively correlated to activity density, *P. cupreus*, *A. dorsalis* and *H. rufipes* while its effect is very variable over other periods (Figure Annexe 4).

The proportion of cultivated area in the landscape accounted for on average 8% of explained variations in carabid distributions (Figure 3 of the main text). Occurrence of *A. similata* was the least response variable influenced by landscape (<1%) while activity density was the most influenced (17%). The proportion of cultivated area always had the same effect on total activity density and species richness with an inverted hump shaped effect. Carabids are the least abundant and diverse in landscapes having a proportion of about 80% and 68% of cultivated area. On the opposite, *P. cupreus* and *H. rufipes* occur the most in landscape composed by 66% and 69% of cultivated area in summer models (Figure of this annexe and Table S3).

‘Region’ explained about 20% of carabid distributions (Figure 2 of the main text and Table S3). Occurence of *A. dorsalis* was the least influenced response by region (3%) and activity density was the most influenced by region (around 30%).

**Figure Annexe 4.**
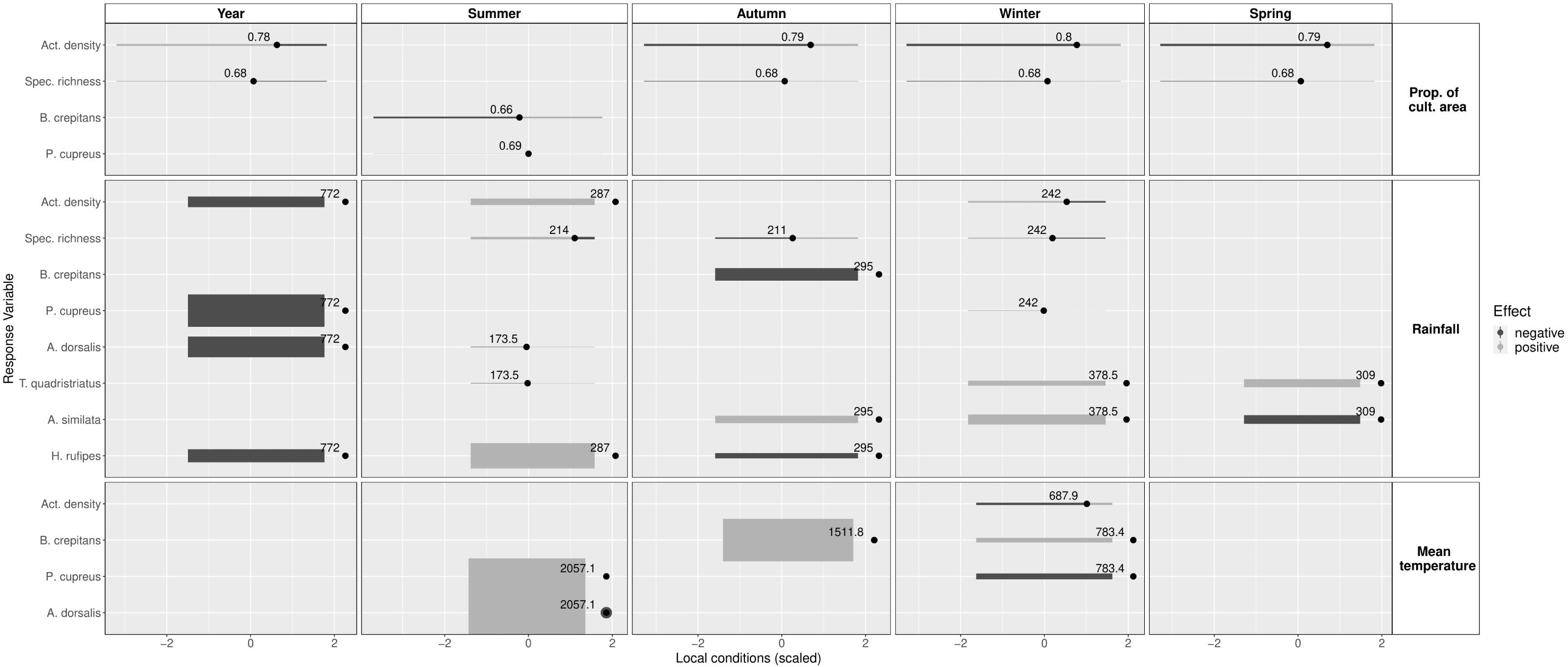
Carabid responses to meteorological conditions and landscape context. Two-tone rights represent quadratic effects with a negative slope for the blue section of the right and a positive slope for the red portion. One-tone right displays simple linear effects. Right thicknesses represent the intensity of the effect, and are proportional to the magnitude the parameter *c* of each predictor. Only the significant effects are represented (See Table S3).

### Supplementary discussion

#### Contribution of the environmental context to explain carabid distributions

In the majority of our models, we found that meteorological conditions all along the year were the most influencing factor driving carabid distribution during the spring-summer period. Mean temperature is the most influential factor in previous summer while rainfall is the most influencing factor during previous autumn, and winter. Rainfall has already been identified as a driver of carabid distribution via change of species composition, not activity density or richness (Skarbek et al., 2021). In this line, evidence shows that soil moisture in winter benefits carabid emergence (Holland et al., 2007), females carabids could prefer oviposition sites with adapted moisture conditions.

Furthermore, the effect of temperature on carabid distribution is contrasted, it has been identified as an important driver of overwintering success in some cases but not in others (Roume et al., 2011), which we did not detect in our study. To our knowledge, no study has evaluated the effect of the previous summer on carabid distribution during the following spring-summer period. However, our results suggest that temperature increase over the 5 year-period is in a range that benefits carabid in agrosystems. Some carabid species lay more eggs of wider size under higher temperature (Ernsting et Isaaks, 2001). Finally, we found very few studies on the effect of meteorological data on carabid distribution in ecosystems (e.g. Jaskuła et Soszyńska-Maj, 2011). We argue further studies on carabids to include meteorological data among their set of predictors to increase the explained variance of their data.

Landscape context, notably landscape composition has been identified as a driver of species richness (e.g. Weibull et al., 2003; Woodcock et al., 2010) but its effect is not consistent among studies (e.g. Puech et al., 2015; Riggi et al., 2017). Here, we showed that according to the range of values of landscape composition, its effects varied, being either negative below 68% to 80% of the proportion of cultivated area or positive above the threshold. These suggest that in complex landscapes, fields could benefit from forest species while dominant (ruderal) species could be highly favored in simple landscapes (Winqvist et al., 2011). This change of effects perceived with the integration of quadratic effects in our models could explain the high variability of carabid responses according to studies (Karp et al., 2018). However, our quadratic effects do not increase the variance explained by landscape composition because we only explained about 8%. Although a few studies quantify the relative contribution of covariates to explained variances, our results are in the same range of values as other studies (e.g. Aviron et al., 2005; Trichard et al., 2013). Additional descriptors describing landscape configuration might increase the effects of landscape on carabid variations (Martin et al., 2019; Carbonne et al., 2022).

## REFERENCES

Aguilera, G., Riggi, L., Miller, K., Roslin, T., & Bommarco, R. (2021). Organic fertilisation enhances generalist predators and suppresses aphid growth in the absence of specialist predators. Journal of Applied Ecology, 58(7), 1455–1465. https://doi.org/10.1111/1365-2664.13862

Allart, R., Ricci, B., & Poggi, S. (2021). Procédure alm de cartographie automatique du paysage. Cahiers Des Techniques de l’INRA, 103, 12.

Barton, K. (2020). MuMIn: Multi-Model Inference. R package version 1.43.17. https://CRAN.R-project.org/package=MuMIn

Bates DM, Maechler M, Bolker B, Walker S (2015). Fitting Linear Mixed-Effects Models Using lme4.” Journal of Statistical Software, 67(1), 148. doi:10.18637/jss.v067.i01.

Bendig, J., Yu, K., Aasen, H., Bolten, A., Bennertz, S., Broscheit, J., Gnyp, M. L., & Bareth, G. (2015). Combining UAV-based plant height from crop surface models, visible, and near infrared vegetation indices for biomass monitoring in barley. International Journal of Applied Earth Observation and Geoinformation, 39, 79–87. https://doi.org/10.1016/j.jag.2015.02.012

Benjamini, Y., Drai, D., Elmer, G., Kafkafi, N., & Golani, I. (2001). Controlling the false discovery rate in behavior genetics research. Behavioural Brain Research, 125(1), 279–284. https://doi.org/10.1016/S0166-4328(01)00297-2

Burnham, K. P., Anderson, D. R., & Huyvaert, K. P. (2011). AIC model selection and multimodel inference in behavioral ecology: Some background, observations, and comparisons. Behavioral Ecology and Sociobiology, 65(1), 23–35. https://doi.org/10.1007/s00265-010-1029-6

Carbonne, B., Bohan, D. A., Foffová, H., Daouti, E., Frei, B., Neidel, V., Saska, P., Skuhrovec, J., & Petit, S. (2022). Direct and indirect effects of landscape and field management intensity on carabids through trophic resources and weeds. Journal of Applied Ecology, 59, 176–187. https://doi.org/10.1111/1365-2664.14043

Climatik database, INRAE, https://agroclim.inrae.fr/climatik/ClimatikGwt.html

Damour, G., Navas, M. L., & Garnier, E. (2018). A revised trait-based framework for agroecosystems including decision rules. Journal of Applied Ecology, 55(1), 12–24.

Dennis, P., Thomas, M. B., & Sotherton, N. W. (1994). Structural Features of Field Boundaries Which Influence the Overwintering Densities of Beneficial Arthropod Predators. Journal of Applied Ecology, 31(2), 361–370. https://doi.org/10.2307/2404550

Diekötter, T., Wamser, S., Wolters, V., & Birkhofer, K. (2010). Landscape and management effects on structure and function of soil arthropod communities in winter wheat. Agriculture, Ecosystems & Environment, 137(1), 108–112. https://doi.org/10.1016/j.agee.2010.01.008

Djoudi, E. A., Marie, A., Mangenot, A., Puech, C., Aviron, S., Plantegenest, M., & Pétillon, J. (2018). Farming system and landscape characteristics differentially affect two dominant taxa of predatory arthropods. Agriculture, Ecosystems & Environment, 259, 98–110. https://doi.org/10.1016/j.agee.2018.02.031

Ephy website (2016) Le catalogue des produits phytopharmaceutiques et de leurs usages des matières fertilisantes et des Supports de culture homologués en France. https://ephy.anses.fr/.

Eyre, M. D., Luff, M. L., & Leifert, C. (2013). Crop, field boundary, productivity and disturbance influences on ground beetles (Coleoptera, Carabidae) in the agroecosystem. Agriculture, Ecosystems & Environment, 165, 60–67. https://doi.org/10.1016/j.agee.2012.12.009

Eyre, M. D., Rushton, S. P., Luff, M. L., & Telfer, M. G. (2005) Investigating the relationships between the distribution of British ground beetle species (Coleoptera, Carabidae) and temperature, precipitation and altitude. Journal of Biogeography, 32(6), 973–983. https://doi.org/10.1111/j.1365-2699.2005.01258.x

Fernández-Tizón, M., Emmenegger, T., Perner, J., & Hahn, S. (2020). Arthropod biomass increase in spring correlates with NDVI in grassland habitat. The Science of Nature, 107(5), 42. https://doi.org/10.1007/s00114-020-01698-7

Frei, B., Guenay, Y., Bohan, D. A., Traugott, M., & Wallinger, C. (2019). Molecular analysis indicates high levels of carabid weed seed consumption in cereal fields across Central Europe. Journal of Pest Science, 92(3), 935–942.

Gaba, S., Fried, G., Kazakou, E., Chauvel, B., & Navas, M.-L. (2014). Agroecological weed control using a functional approach: A review of cropping systems diversity. Agronomy for Sustainable Development, 34(1), 103–119.

Gagic, V., Kleijn, D., Báldi, A., Boros, G., Jørgensen, H. B., Elek, Z., Garratt, M. P., de Groot, G. A., Hedlund, K., & Kovács-Hostyánszki, A. (2017). Combined effects of agrochemicals and ecosystem services on crop yield across Europe. Ecology Letters, 20(11), 1427–1436.

Geiger, F., Bengtsson, J., Berendse, F., Weisser, W. W., Emmerson, M., Morales, M. B., Ceryngier, P., Liira, J., Tscharntke, T., Winqvist, C., Eggers, S., Bommarco, R., Pärt, T., Bretagnolle, V., Plantegenest, M., Clement, L. W., Dennis, C., Palmer, C., Oñate, J. J., … Inchausti, P. (2010). Persistent negative effects of pesticides on biodiversity and biological control potential on European farmland. Basic and Applied Ecology, 11(2), 97–105. https://doi.org/10.1016/j.baae.2009.12.001

Hartig, F. (2021). DHARMa: Residual Diagnostics for Hierarchical (Multi-Level / Mixed) Regression Models. R package version 0.4.1. https://CRAN.R-project.org/package=DHARMa

Henneron, L., Bernard, L., Hedde, M., Pelosi, C., Villenave, C., Chenu, C., Bertrand, M., Girardin, C., & Blanchart, E. (2015). Fourteen years of evidence for positive effects of conservation agriculture and organic farming on soil life. Agronomy for Sustainable Development, 35(1), 169–181.

Honek, A. (1997). The effect of temperature on the activity of Carabidae (Coloptera) in a fallow field. Eur. J. Entomol., 94 (1997), pp. 97–104.

Jowett, K., Milne, A. E., Garrett, D., Potts, S. G., Senapathi, D., & Storkey, J. (2021). Above- and below-ground assessment of carabid community responses to crop type and tillage. Agricultural and Forest Entomology, 23(1), 1–12. https://doi.org/10.1111/afe.12397

Jowett, K., Milne, A. E., Metcalfe, H., Hassall, K. L., Potts, S. G., Senapathi, D., & Storkey, J. (2019). Species matter when considering landscape effects on carabid distributions. Agriculture, Ecosystems & Environment, 285, 106631. https://doi.org/10.1016/j.agee.2019.106631

Karp, D. S., Chaplin-Kramer, R., Meehan, T. D., Martin, E. A., DeClerck, F., Grab, H., Gratton, C., Hunt, L., Larsen, A. E., Martínez-Salinas, A., O’Rourke, M. E., Rusch, A., Poveda, K., Jonsson, M., Rosenheim, J. A., Schellhorn, N. A., Tscharntke, T., Wratten, S. D., Zhang, W., … Zou, Y. (2018). Crop pests and predators exhibit inconsistent responses to surrounding landscape composition. Proceedings of the National Academy of Sciences, 115(33), E7863–E7870. https://doi.org/10.1073/pnas.1800042115

Kleyer, M., Bekker, R. M., Knevel, I. C., Bakker, J. P., Thompson, K., Sonnenschein, M., Poschlod, P., Groenendael, J. M. V., Klimeš, L., Klimešová, J., Klotz, S., Rusch, G. M., Hermy, M., Adriaens, D., Boedeltje, G., Bossuyt, B., Dannemann, A., Endels, P., Götzenberger, L., … Peco, B. (2008). The LEDA Traitbase: A database of life-history traits of the Northwest European flora. Journal of Ecology, 96(6), 1266–1274. https://doi.org/10.1111/j.1365-2745.2008.01430.x

Labruyere, S., Ricci, B., Lubac, A., & Petit, S. (2016). Crop type, crop management and grass margins affect the abundance and the nutritional state of seed-eating carabid species in arable landscapes. Agriculture, Ecosystems & Environment, 231, 183–192. https://doi.org/10.1016/j.agee.2016.06.037

Mahaut, L., Gaba, S., & Fried, G. (2019). A functional diversity approach of crop sequences reveals that weed diversity and abundance show different responses to environmental variability. Journal of Applied Ecology, 56(6), 1400–1409.

Marrec, R., Caro, G., Miguet, P., Badenhausser, I., Plantegenest, M., Vialatte, A., Bretagnolle, V., & Gauffre, B. (2017). Spatiotemporal dynamics of the agricultural landscape mosaic drives distribution and abundance of dominant carabid beetles. Landscape Ecology, 32(12), 2383–2398. https://doi.org/10.1007/s10980-017-0576-x

Marrec, R., Gross, N., Badenhausser, I., Dupeyron, A., Caro, G., Bretagnolle, V., Roncoroni, M., & Gauffre, B. (2021). Functional traits of carabid beetles reveal seasonal variation in community assembly in annual crops (p. 2021.02.04.429696). https://doi.org/10.1101/2021.02.04.429696

Petit, S., Deytieux, V., & Cordeau, S. (2021). Landscape-scale approaches for enhancing biological pest control in agricultural systems. Environmental Monitoring and Assessment, 193(1), 1–13.

Petit, S., Muneret, L., Carbonne, B., Hannachi, M., Ricci, B., Rusch, A., & Lavigne, C. (2020). Chapter One - Landscape-scale expansion of agroecology to enhance natural pest control: A systematic review. In D. A. Bohan & A. J. Vanbergen (Eds.), Advances in Ecological Research (Vol. 63, pp. 1–48). Academic Press. https://doi.org/10.1016/bs.aecr.2020.09.001

Purtauf, T., Dauber, J., & Wolters, V. (2005). The response of carabids to landscape simplification differs between trophic groups. Oecologia, 142(3), 458–464. https://doi.org/10.1007/s00442-004-1740-y

R Core Team (2021). R: A language and environment for statistical computing. R Foundation for Statistical Computing, Vienna, Austria. https://www.R-project.org/

Redlich, S., Martin, E. A., & Steffan-Dewenter, I. (2018). Landscape-level crop diversity benefits biological pest control. Journal of Applied Ecology, 55(5), 2419–2428. https://doi.org/10.1111/1365-2664.13126

Riggi, L. G. A., & Bommarco, R. (2019). Subsidy type and quality determine direction and strength of trophic cascades in arthropod food webs in agroecosystems. Journal of Applied Ecology, 56(8), 1982–1991. https://doi.org/10.1111/1365-2664.13444

Roubinet, E., Jonsson, T., Malsher, G., Staudacher, K., Traugott, M., Ekbom, B., & Jonsson, M. (2018). High Redundancy as well as Complementary Prey Choice Characterize Generalist Predator Food Webs in Agroecosystems. Scientific Reports, 8(1), 8054. https://doi.org/10.1038/s41598-018-26191-0

Schellhorn, N. A., Gagic, V., & Bommarco, R. (2015). Time will tell: Resource continuity bolsters ecosystem services. Trends in Ecology & Evolution, 30(9), 524–530.

Shearin, A. F., Reberg-Horton, S. C., & Gallandt, E. R. (2008). Cover Crop Effects on the Activity-Density of the Weed Seed Predator Harpalus rufipes (Coleoptera: Carabidae). Weed Science, 56(3), 442–450. https://doi.org/10.1614/WS-07-137.1

Shennan, C., Krupnik, T. J., Baird, G., Cohen, H., Forbush, K., Lovell, R. J., & Olimpi, E. M. (2017). Organic and conventional agriculture: A useful framing? Annual Review of Environment and Resources, 42, 317–346.

Tamburini, G., Bommarco, R., Wanger, T. C., Kremen, C., van der Heijden, M. G. A., Liebman, M., & Hallin, S. (2020). Agricultural diversification promotes multiple ecosystem services without compromising yield. Science Advances, (6), eaba1715. https://doi.org/10.1126/sciadv.aba1715

Timberlake, T. P., Vaughan, I. P., & Memmott, J. (2019). Phenology of farmland floral resources reveals seasonal gaps in nectar availability for bumblebees. Journal of Applied Ecology, 56(7), 1585–1596. https://doi.org/10.1111/1365-2664.13403

Uhler, J., Redlich, S., Zhang, J., Hothorn, T., Tobisch, C., Ewald, J., Thorn, S., Seibold, S., Mitesser, O., Morinière, J., Bozicevic, V., Benjamin, C. S., Englmeier, J., Fricke, U., Ganuza, C., Haensel, M., Riebl, R., Rojas-Botero, S., Rummler, T., … Müller, J. (2021). Relationship of insect biomass and richness with land use along a climate gradient. Nature Communications, 12(1), 5946. https://doi.org/10.1038/s41467-021-26181-3

Vasseur, C., Joannon, A., Aviron, S., Burel, F., Meynard, J.-M., & Baudry, J. (2013). The cropping systems mosaic: How does the hidden heterogeneity of agricultural landscapes drive arthropod populations? Agriculture, Ecosystems & Environment, 166, 3–14.

Ward, M. J., Ryan, M. R., Curran, W. S., Barbercheck, M. E., & Mortensen, D. A. (2011). Cover Crops and Disturbance Influence Activity-Density of Weed Seed Predators Amara aenea and Harpalus pensylvanicus (Coleoptera: Carabidae). Weed Science, 59(1), 76–81. https://doi.org/10.1614/WS-D-10-00065.1

Winqvist, C., Bengtsson, J., Aavik, T., Berendse, F., Clement, L. W., Eggers, S., Fischer, C., Flohre, A., Geiger, F., Liira, J., Pärt, T., Thies, C., Tscharntke, T., Weisser, W. W., & Bommarco, R. (2011). Mixed effects of organic farming and landscape complexity on farmland biodiversity and biological control potential across Europe. Journal of Applied Ecology, 48(3), 570–579. https://doi.org/10.1111/j.1365-2664.2010.01950.x

Yin, X., Hayes, R. M., McClure, M. A., & Savoy, H. J. (2012). Assessment of plant biomass and nitrogen nutrition with plant height in early-to mid-season corn. Journal of the Science of Food and Agriculture, 92(13), 2611–2617. https://doi.org/10.1002/jsfa.5700

## Supplementary references

Aviron, S., Burel, F., Baudry, J., & Schermann, N. (2005). Carabid assemblages in agricultural landscapes: Impacts of habitat features, landscape context at different spatial scales and farming intensity. Agriculture, Ecosystems & Environment, 108(3), 205–217. https://doi.org/10.1016/j.agee.2005.02.004

Carbonne, B., Bohan, D. A., Foffová, H., Daouti, E., Frei, B., Neidel, V., Saska, P., Skuhrovec, J., & Petit, S. (2022). Direct and indirect effects of landscape and field management intensity on carabids through trophic resources and weeds. Journal of Applied Ecology, 59, 176–187. DOI: 10.1111/1365-2664.14043

Ernsting, G., & Isaaks, A. (2000). Ectotherms, Temperature, and Trade-offs: Size and Number of Eggs in a Carabid Beetle. The American Naturalist, 155(6), 804–813. https://doi.org/10.1086/303361

Holland, J. M., Thomas, C. F. G., Birkett, T., & Southway, S. (2007). Spatio-temporal distribution and emergence of beetles in arable fields in relation to soil moisture. Bulletin of Entomological Research, 97(1), 89–100. https://doi.org/10.1017/S0007485307004804

Jaskula, R., & Soszyńska-Maj, A. (2011). What do we know about winter active ground beetles (Coleoptera, Carabidae) in Central and Northern Europe? ZooKeys, 100, 517–532. https://doi.org/10.3897/zookeys.100.1543

Martin, E. A., Dainese, M., Clough, Y., Báldi, A., Bommarco, R., Gagic, V., Garratt, M. P. D., Holzschuh, A., Kleijn, D., Kovács-Hostyánszki, A., Marini, L., Potts, S. G., Smith, H. G., Al Hassan, D., Albrecht, M., Andersson, G. K. S., Asís, J. D., Aviron, S., Balzan, M. V., … Steffan-Dewenter, I. (2019). The interplay of landscape composition and configuration: New pathways to manage functional biodiversity and agroecosystem services across Europe. Ecology Letters, 22(7), 1083–1094. https://doi.org/10.1111/ele.13265

Puech, C., Poggi, S., Baudry, J., & Aviron, S. (2015). Do farming practices affect natural enemies at the landscape scale? Landscape Ecology, 30(1), 125–140. https://doi.org/10.1007/s10980-014-0103-2

Riggi, L. G., Gagic, V., Rusch, A., Malsher, G., Ekbom, B., & Bommarco, R. (2017). Pollen beetle mortality is increased by ground-dwelling generalist predators but not landscape complexity. Agriculture, Ecosystems & Environment, 250, 133–142. https://doi.org/10.1016/j.agee.2017.06.039

Roume A., 2011. Quelle est la contribution des milieux semi naturels à la diversité et à la répartition des assemblages de Carabidae circulants et hivernants dans un paysage rural tempéré ? https://www.theses.fr/2011INPT0043

Skarbek, C. J., Kobel-Lamparski, A., & Dormann, C. F. (2021). Trends in monthly abundance and species richness of carabids over 33 years at the Kaiserstuhl, southwest Germany. Basic and Applied Ecology, 50, 107–118. https://doi.org/10.1016/j.baae.2020.11.003

Trichard, A., Alignier, A., Biju-Duval, L., & Petit, S. (2013). The relative effects of local management and landscape context on weed seed predation and carabid functional groups. Basic and Applied Ecology, 14(3), 235–245.

Weibull, A.-C., & Östman, Ö. (2003). Species composition in agroecosystems: The effect of landscape, habitat, and farm management. Basic and Applied Ecology, 4(4), 349–361. https://doi.org/10.1078/1439-1791-00173

Woodcock, B. A., Redhead, J., Vanbergen, A. J., Hulmes, L., Hulmes, S., Peyton, J., Nowakowski, M., Pywell, R. F., & Heard, M. S. (2010). Impact of habitat type and landscape structure on biomass, species richness and functional diversity of ground beetles. Agriculture, Ecosystems & Environment, 139(1), 181–186. https://doi.org/10.1016/j.agee.2010.07.018

